# The TaxUMAP atlas: efficient display of large clinical microbiome data reveals ecological competition involved in protection against bacteremia

**DOI:** 10.1101/2022.09.27.509746

**Authors:** Jonas Schluter, Ana Djukovic, Bradford P. Taylor, Jinyuan Yan, Caichen Duan, Grant A. Hussey, Chen Liao, Sneh Sharma, Emily Fontana, Luigi A. Amoretti, Roberta J. Wright, Anqi Dai, Jonathan U. Peled, Ying Taur, Miguel-Angel Perales, Benjamin A. Siranosian, Ami S. Bhatt, Marcel R.M. van den Brink, Eric G. Pamer, Joao B. Xavier

## Abstract

The microbiome is associated with health and disease, but causal effects are hard to quantify— especially in humans where controlled experiments are nearly impossible. Akin to natural experiments, closely monitored patients offer an alternative to characterize microbiome effects. We present TaxUMAP, a taxonomically-informed visualization method to effectively display diverse microbiome states. TaxUMAP charts a microbiome atlas from 1,870 cancer patients as they progress through therapy-induced perturbations, and quantifies the microbiome contribution to patients’ risk for life-threatening bacteremia. We find that the lowest diversity states (gut dominations) that follow antibiotic treatments are stable, and that diverse communities harbor more diverse antimicrobial resistance genes than dominations. We reveal that certain *Klebsiella* species are associated with reduced risk for bacteremia, an effect driven by bacterial competition that we validate experimentally *in vitro* and *in vivo*. TaxUMAP effectively maps longitudinal microbiome data that can facilitate research into causal microbiome effects on human health.

**HIGHLIGHTS:**

- TaxUMAP charts an atlas of patients’ microbiome states and their clinical context to reveal new causal effects.
- Antibiotics deplete the biodiversity and reduce the number of different antimicrobial resistance genes in the gut microbiome.
- Certain *Klebsiella* species are associated with lower risk of bacteremia by other gut-borne pathogens.
- These *Klebsiella* outcompete other gram-negative pathogens *in vivo*.

## INTRODUCTION

The gut microbiome contains the most numerous and diverse bacterial community in the human body. Some species of gut bacteria have been associated with human health (Manzo and Bhatt, 2015; Olin et al., 2018), for example by reducing the risk of gut colonization by incoming pathogens (Buffie and Pamer, 2013; Caballero et al., 2017). Antibiotics perturb the microbiota composition (Dethlefsen and Relman, 2011), reduce the density of strict anaerobes associated with resistance to pathogen colonization (Buffie et al., 2015; Morjaria et al., 2019) and lead to the expansion of antibiotic-resistant bacteria that increase the risk of bacteremia in immune-compromised patients (Stoma et al., 2021; Taur et al., 2012). At Memorial Sloan Kettering Cancer Center (MSKCC), 50% of the patients hospitalized for allogeneic hematopoietic stem cell transplantation (HCT) have intestinal expansions (dominations) by vancomycin resistant *Enterococcus* (VRE) after antibiotics. This domination increases the risk of VRE bacteremia 9-fold (Taur et al., 2012). Another ∼15% of patients become dominated by gram-negative proteobacteria such as *Escherichia coli* and *Klebsiella pneumoniae*. Those dominations increase the risk of gram-negative bacteremia by >10-fold (Stoma et al., 2021). Similar patterns occur in other transplant centers across the globe (Peled et al., 2020), and in other hospitalized patient populations such as COVID-19 patients (Venzon et al., 2022).

The dominations of the gut microbiome that happen after antibiotics are strongly associated with higher risks of bacteremia in cancer patients receiving systemic chemotherapy (Stoma et al., 2021; Tamburini et al., 2018; Taur et al., 2012). But establishing causal effects for even such strong clinical associations remains difficult. Finding the mechanisms underlying other patient outcomes, such as cancer progression and response to therapy where any association with microbiome composition is perhaps indirect, may be even harder (Xavier et al., 2020). Experimental tests of cause and effect relationships between the microbiome and host phenotypes are easier in animal models; however, due to different ecological and host environments among other factors, the experimental findings do not always translate to humans (Walter et al., 2020). Large longitudinal datasets of patient microbiomes provide an alternative: when the patients experience microbiome compositional changes in response to treatments such as antibiotics, their longitudinal data could be mined to infer relationships of cause and effect (Gerber, 2014; Schluter et al., 2020). This is conceptually similar to how “natural experiments”, events or policy changes that happen in real life, can enable economists to quantify causal effects in human societies (Card and Krueger, 1993).

Here we introduce a new technique to study large scale microbiome data in the context of clinical and host metadata. The method, called TaxUMAP, extends the Uniform Manifold Approximation and Projection (UMAP) to include hierarchical taxonomic relations between the bacterial sequencing variants identified in a sample (Kobak and Linderman, 2021; McInnes et al., 2018) to cluster microbiome community samples with similar global ecosystem features while retaining fine-grained differences. We use TaxUMAP to chart an atlas of microbiome compositions using >10,000 samples from 1,870 patients hospitalized at Memorial Sloan Kettering to receive allo-HCT (Liao et al., 2021; Yan et al., 2022). TaxUMAP enables the hypothesis of new cause-effect relationships suited for experimental validation. We illustrate this use by exploring microbiome states associated with high risk of bloodstream infections. Our study revealed a region of the atlas enriched for certain species of the genus *Klebsiella* excluding other, high infection risk-associated enterobacteria. This observation was intriguing: independent studies have shown that the same types of *Klebsiella* are important for resistance to pathogen colonization in mice (Oliveira et al., 2020; Osbelt et al., 2021). We then isolated strains from corresponding patient samples and experimentally validated their ability to outcompete pathogenic isolates. TaxUMAP facilitates the identification of potential ecological mechanisms in the gut microbiome that have a causal health effect on the host.

### RESULTS

### TaxUMAP charts an atlas from longitudinal clinical microbiota data

We built TaxUMAP to facilitate the development of causal inference of the microbiome effect on human health from a large longitudinal microbiome data set obtained from hospitalized cancer patients (Liao et al., 2021). The immunocompromised state of cancer patients and the many microbiome perturbations they incur could exacerbate health effects of bacterial populations, and therefore make them more visible (Schluter et al., 2020). HCT is a complex procedure with three phases: pre-transplant chemotherapeutic conditioning (phase I), a prolonged period of low immune cell counts (neutropenia, phase II), and immune reconstitution (phase III, **Figure 1A**). The duration of each phase varies among patients; we split samples in each phase evenly into early, mid and late-phase, providing a comparable pseudo time point (**Figure 1B-D**). Anti-infective drugs are administered to patients prophylactically during phase I and II and as needed during any phase for patients with suspected infections. Antibiotics cause collateral damage to the gut ecosystem (Blaser, 2016; Dethlefsen and Relman, 2011; Maier et al., 2021; Morjaria et al., 2019; Niehus et al., 2020; Venzon et al., 2022): microbiota α-diversity tends to drop from phase I to phase II, and to stay low in phase III (**Figure 1C,D**) (Peled et al., 2020).

**Figure 1.**
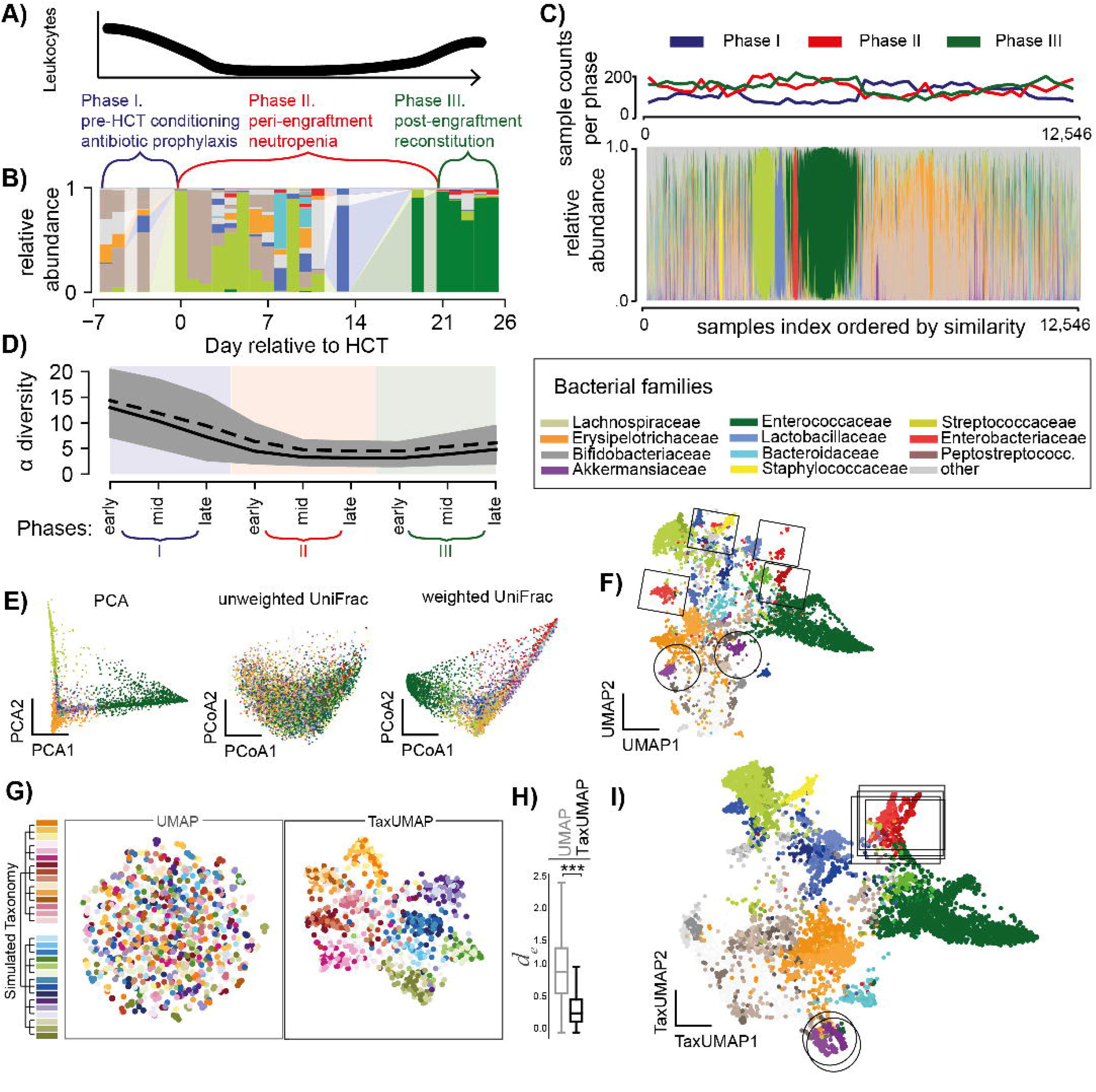
The TaxUMAP algorithm effectively visualizes different microbiome states. A) Three distinct clinical phases of a HCT therapy. B) Gut microbiome bacterial compositions over time from one patient; bars represent the relative taxon abundances measured in a stool sample by 16S rRNA gene sequencing. C) Compositions in 12,546 samples; clinical phases at collection indicated. D) Bacterial alpha diversity (inverse Simpson index) over pseudo-time by assigning each sample to an early, mid, or late phase of the respective clinical phase (shading as per colors in A) during which a sample was obtained; mean (black solid) and median (black dashed) diversity (n=12,546, shaded: 95%C.I. of the mean). E) ASV level principal component, and principal coordinate plots of all samples (unweighted- and weighted UniFrac distances). F) UMAP embedding at the ASV level. G) Comparison between UMAP and TaxUMAP on a simulated data set; colors indicate the hypothetical genus, indicated on the simulated taxonomy tree, with the highest abundance in a sample. H) Euclidean distances between *in silico* generated samples with the same dominant family in UMAP vs. TaxUMAP embedding (n=1,000, ***:p<10^−4^, Wilcoxon rank sum test). I) TaxUMAP embedding of patient samples. E, F, I) color by most abundant taxon.

The microbiota contains hundreds of bacterial taxa. An ideal way to map the microbiome states effectively should reduce the data dimensionality without losing crucial detail. Common ordination methods such as principal component analysis (PCA), or principal coordinate analyses using UniFrac distances (PCoA) can be applied to allo-HCT patient data, but they only resolve the most frequently observed, low diversity, single-taxon enriched microbiota states (**Figure 1E**) at the expense of many other distinct states of microbiota composition that become less visible. The recently developed nonlinear dimensionality reduction method UMAP (McInnes et al., 2018a) is especially useful to work with large sample numbers and high data dimensionality (Armstrong et al., 2021) and reveal microbiota states left unclear in other methods (**Figure 1F**). However, it is missing crucial biological similarity information in the form of taxonomic relationships between ASVs. To demonstrate this, we simulated 1,000 bacterial compositions from 600 AVSs belonging to 30 different hypothetical species, ten families, and 2 phyla (**Figure 1G**). While the UMAP places samples with similar species composition together, it neglects similarities between samples based on high taxonomic levels (**Figure 1F, G**). When applying UMAP to our patient samples at the ASV level, for example, samples dominated by different ASVs from the genus *Akkermansia* (in shades of purple, highlighted by circles, **Figure 1F**), or the family *Enterobacteriaceae* (in shades of red, highlighted by rectangles, **Figure 1F)** end up far from each other.

Our open-source TaxUMAP algorithm combines ecological distance matrices computed at different taxonomy levels. Each taxonomic level can be emphasized or disregarded by the user, depending on the research question. The resulting matrix of taxonomically aware pairwise sample distances is then used as input for the UMAP. When applying this technique to our in silico data (**Figure 1G**), formerly separate clusters of samples with different species but similar families are now embedded significantly closer together (**Figure 1H**). In our patient data, the highlighted clusters that were artefactually far apart (**Figure 1F**) are now moved together (**Figure 1I**), facilitating a biologically interpretable visualization.

### Atlas annotation with clinical data reveals collateral damage caused by antibiotics

The TaxUMAP atlas charts clinical metadata onto microbiome states. While the dataset has slightly more men than women, the atlas reveals that sex does not play an important role, since patients of both sexes visit the same regions (**Figure 2A**). Nor does the underlying disease (**Figure 2B**). Every patient, independent of sex or disease, received antibiotic prophylaxis. Phase I samples fall in the bottom left corner of the map (**Figure 2C**), with the earliest samples (**Figure 2D**) preceding anti-infective treatment (**Figure 2E**). Antibiotics precede changes in the gut microbiome composition, particularly a loss of diversity (**Figure 2F**) and reduction in the relative abundances of obligate anaerobes (**Figure 2G, S1**). Alongside these high-level shifts in the microbiome, the consistency of stool also changes from well-formed to semi-formed and liquid (**Figure 2H**). The total bacterial density (16S rRNA gene copies per gram of stool) varies widely, from ∼10^10^ copies per gram of stool, to almost undetectable in some samples (**Figure 2I, S2A**, limit of detection: 2.84 copies/g stool). Bacterial density correlates positively with community diversity in non-liquid samples but negatively in liquid samples (**Figure S2B**). Microbiome compositional data alone suffers from a lack of total population density information; comparing microbiota composition (**Figure 1I**) with bacterial loads (**Figure 2I**) indicates that ecosystems where Streptococacceae or Staphylococcaceae make up the most frequent bacteria tend to have the lowest densities. Similarly low-diverse ecosystems populated predominantly by Enterobacteriaceae or Enterococcaceae also exhibit low population densities in many samples; however, they also include samples with high bacterial loads, indicating that these bacteria can expand in numbers in the gut during periods where they dominate the community (**Figure 1I, 2I**).

**Figure 2.**
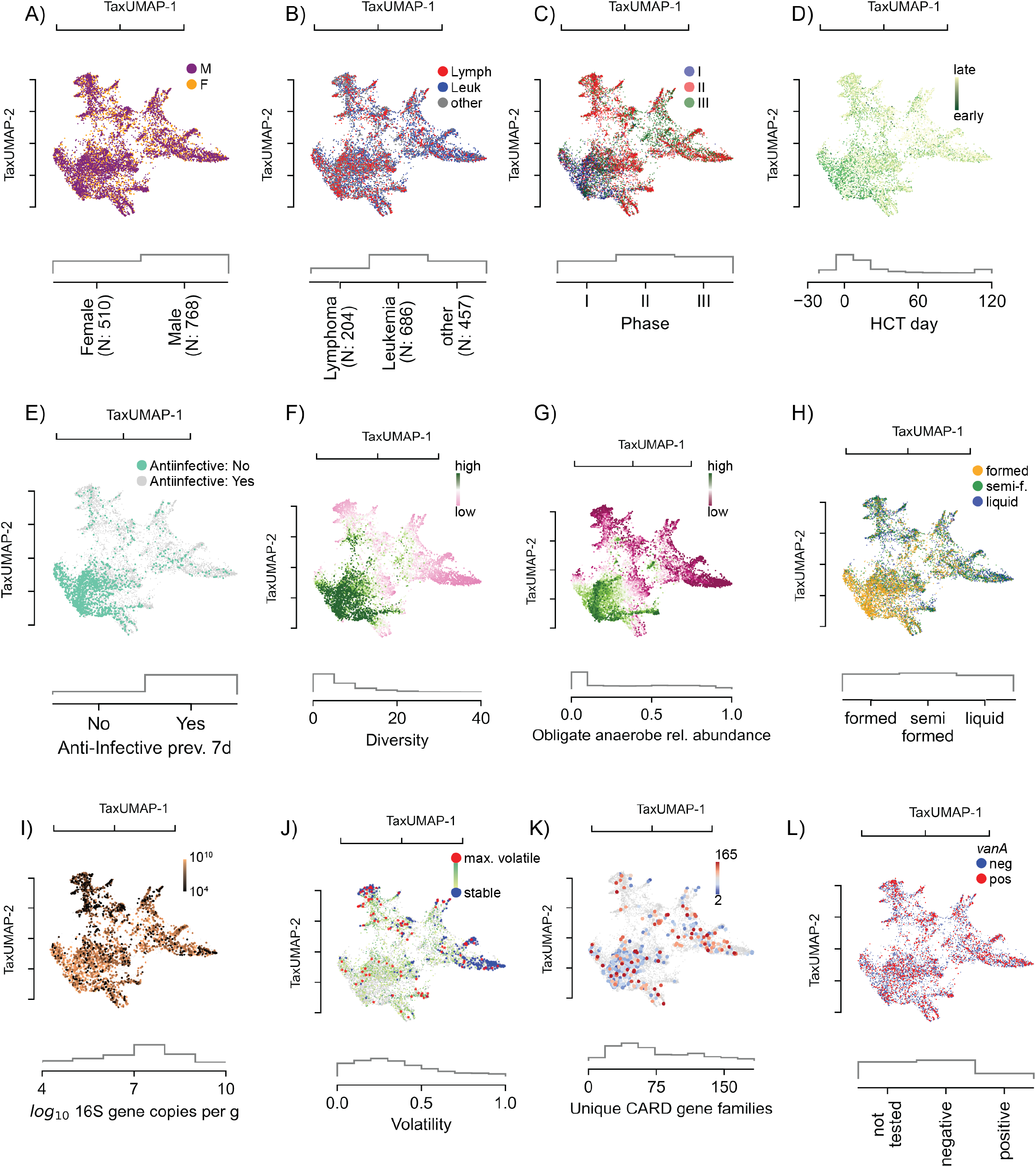
The atlas of the HCT bacterial gut microbiota using TaxUMAP. Each scatter plot is accompanied by histograms (grey) of the displayed metadata distribution. A) Samples from female and male donors, with B) different diseases. C) Samples from different clinical phases; phase I samples are concentrated in a distinct region corresponding to D) samples taken early during therapy. In the same region, samples are concentrated where no anti-infective administrations were recorded in the 7 days prior to sample collection (E). F) Bacterial alpha-diversity (measured by the inverse Simpson index). G) Relative abundances of obligate anaerobe commensal taxa. See also **Figure S1**. H) Stool composition (liquid, semi-formed, formed stool). I) Total bacterial abundance estimated by total 16S gene copy numbers per gram of stool. See also **Figure S2**. J) Volatility of the bacterial community (most volatile in red: volatility >0.9; least volatile in blue: volatility<0.1). See also **Figure S3, S4**. K) Unique antimicrobial resistance phenotypes detected per sample. See also **Figure S5, S6, S7**. L) Vancomycin-resistance conveying *vanA* gene detected in rectal swab. See also **Figure S8**.

The high temporal frequency of the longitudinal data allows us to analyze consecutive time points from the same patient and assess compositional volatility (**Figure 2J**). We could find volatile switches, defined as more than 90% community turnover between two consecutive days, throughout the entire atlas (**Figure S3**), indicating that any microbiome state in allo-HCT patients is prone to a sudden change. Interestingly, the endpoints of volatile switches, which tended to be localized in low-diversity ecosystems, also harbored some of the most stable states, *i*.*e*. where volatility scores were the lowest (**Figure 2J, blue)**. This suggests that the microbiota, once injured, may remain depleted, possibly due to continued selection pressure, for example by antibiotics (Morjaria et al., 2019; Niehus et al., 2020) or because, according to ecological theory, less diverse microbiome ecosystems tend to be more stable (Coyte et al., 2015; May, 1972).

To test if microbiome domination states persist due to continuous antibiotic pressure, we analyzed antibiotic treatment records and correlated them to microbiome composition and diversity (**Figure S4A**). The data showed that several low diversity intestinal domination states can indeed persist beyond the window of antibiotic administration. Biodiversity recovers slowly after the last day of antibiotic treatment, with an average doubling time of >40 days (**Figure S4B**). We quantified the half-life for eight of the most frequent domination states post-antibiotics, and made a comparison among the different taxa (**Figure S4C**). For example, *Enterococcus* (the most frequent) has a half-life of 6.4 days after antibiotics, while *Streptococcus* (the second most frequent) has 15.5 days half-life. While it is possible that antibiotics persist in the gut lumen even after administration was stopped, these results suggest that persistence of dominations might at least in part be due to the nature of the dominating species or the interactions between the rest of the microbiome, including the host, with the dominating species. This is, to our knowledge, the first quantification of the stability of intestinal dominations done directly in patients.

To understand how antibiotics cause the shifts in microbiome composition we analyzed shotgun metagenomic sequences from 395 samples from 49 unique patients. We counted the number of unique, qualitative antibiotic resistance genes (ARG) found in each sample, as done before (Montassier et al., 2021). Plotting the unique ARGs identified in a sample on the TaxUMAP suggested a correspondence with bacterial alpha diversity (**Figure 2F,K**). We found a more diverse set of antimicrobial resistance genes in ecologically diverse microbial samples than in samples dominated by a single taxon (**Figure 2K, S5**). This was an interesting observation because domination states tend to occur after antibiotics, and they are assumed to be caused by selection pressures benefiting antibiotic resistant taxa. Our results suggest, however, that antibiotics, which deplete ecological diversity, act consistent with a model where they purge antimicrobial resistance genes from the ecological community that do not convey resistance to the applied antibiotic, and thereby these antibiotics may also decrease the diversity of the antimicrobial resistance gene pool.

To show how antibiotics can increase the abundance of a specific antimicrobial-resistance gene, we performed a statistical analysis of the diversity-depleting effects of different antibiotics; similar to previous results (Morjaria et al., 2019), we identified significant associations between piperacillin/tazobactam, metronidazole, meropenem and vancomycin with microbiome diversity loss (see **Figure S6A** for a full list). Then, we investigated which ARGs correlated with the administration of those four antibiotics. As expected the number of genes statistically significantly correlated with the antibiotics was only a fraction of the total number of the ARGs (**Figure S6B**). ARGs significantly associated with resistance to these four antibiotics include *blaZ* (encoding a beta-lactamase) and *mecA* (an alternative penicillin-binding protein) of *S. aureus* a taxon that in our data increased in abundance with the administration of the beta-lactam antibiotic piperacillin/tazobactam, as well as several genes of the *vanA* operon associated with vancomycin resistance in *E. faecium*, a taxon that increased in abundance with the administration of oral vancomycin. We detected *vanA* in 54.5% of our shotgun-sequenced samples, all of which showed a sequence identity of >98% with the matched *vanA* gene of *E. faecium* (**Figure S7**). Vancomycin is among the most frequently administered antibiotics in these patients (24.6% of all administrations). We detected the presence of *vanA* in 7,589 samples using a PCR test (**Figure S8A**) localized across the TaxUMAP, but more frequently in the low diversity samples dominated by *Enterococcus* (**Figure 2L**). This agrees with the notion that strong selection for vancomycin resistance, facilitated by the antibiotic-induced loss of obligate anaerobe bacteria, which might otherwise hinder its expansion (Caballero et al., 2017; Ubeda et al., 2010, 2013), drives enterococcal domination. In support of this idea, the patients who were *vanA* positive already in phase I, i.e. prior to vancomycin prophylaxis, tended to lose more diversity in phases II and III (**Figure S8B**) and become dominated by *Enterococcus* (**Figure S8B**).

### Risk of bloodstream infection changes with microbiota ecology

To investigate how strong selection and gut microbiota ecology affect patients’ health, we turned to what is perhaps the most immediate impact of altered microbiome community in these patients: translocation of gut bacteria into the blood leading to bacteremia (Whangbo et al., 2017). Previous studies have linked gut dominations by *Enterococcus* and gram negative proteobacteria to bacteremia (Stoma et al., 2021; Taur et al., 2012). For our analyses, we therefore selected the last stool sample collected ⩽7 days before a patient developed an enterococcal or gamma proteobacterial bacteremia (**Figure 3A**). This showed—consistent with previous studies—that most enterococcal infections followed Enterococcaceae domination and most gamma proteobacterial infections followed gamma proteobacteria domination. To identify specific ASVs associated with infection, we compiled a list of 10 candidate ASVs that were either the most abundant or had the largest increase prior to infections (see methods). BLAST analysis revealed that their 16S rRNA gene sequences matched the bloodstream isolate; each of these 10 ASVs are therefore a potential risk factor for gut-to-blood translocation, especially if they were enriched in patients who later had a BSI relative to similar patients who did not have a BSI. To test this statistically, we conducted a multivariate Bayesian logistic regression comparing BSI-cases to matched samples from uninfected patients (methods); the model takes as input the abundances of all 10 candidate ASVs in a stool sample and predicts the infection risk (AUC: 0.73, **Figure 3C**). The posterior distributions confirmed that 3 out of those 10 ASVs—one belonging to the Enterococcaceae and two to the gamma proteobacterial Enterobacteriaceae family—were positively associated with a higher risk of infection (**Figure 3D**). We displayed the calculated risk score predicted by the model on the atlas to chart regions of highest infection risk (**Figure 3B**). A region located in the enterobacterial-enriched region (**Figure 3B inset**) had the highest risk. This region is enriched in ASV-3, classified as a group of *Escherichia-Shigella* and further characterized by metagenomic sequencing (**Figure 3E, Figure 4**).

**Figure 3.**
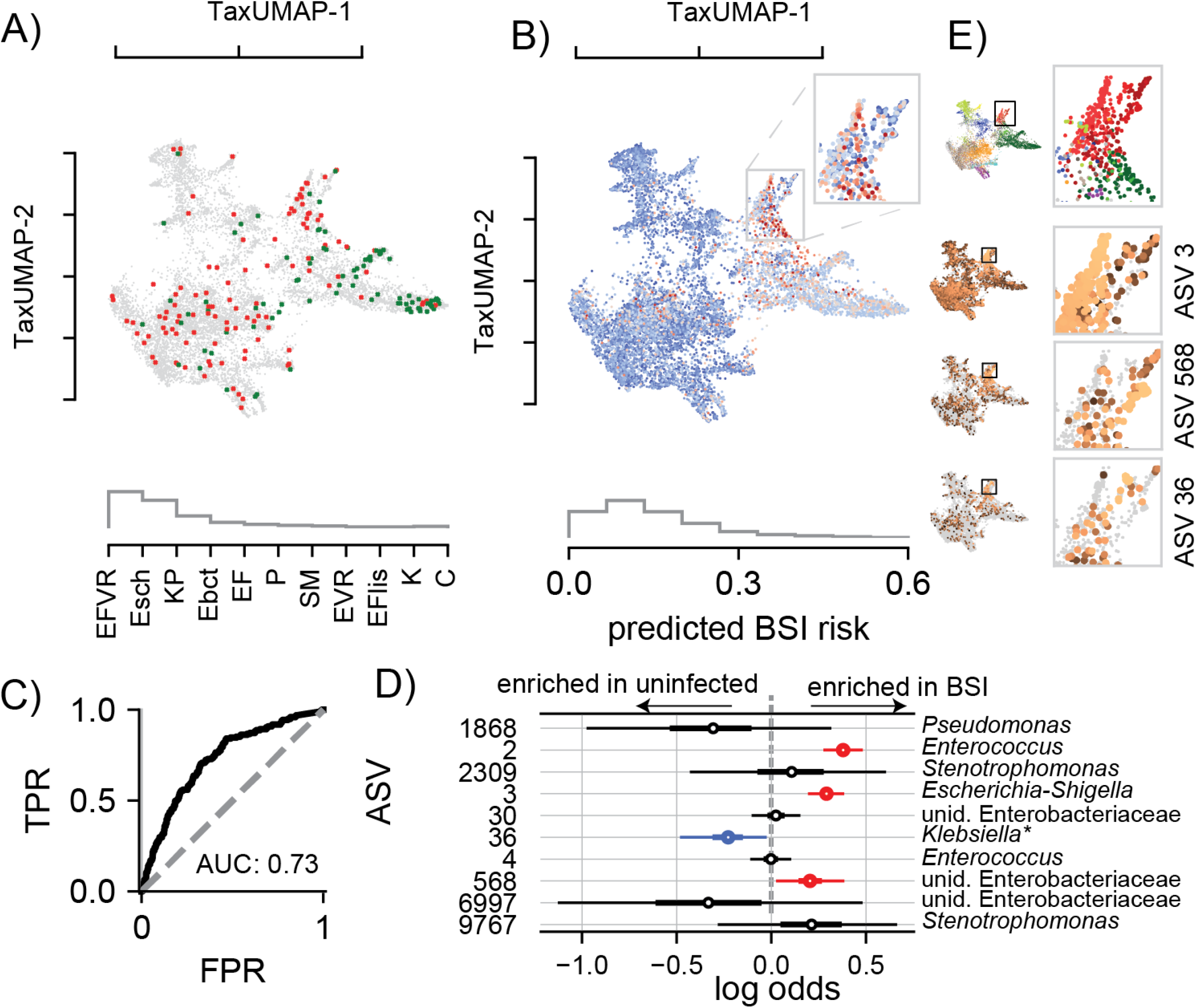
BSI risk scores across the TaxUMAP suggests reduced risk-associated ASV-36-*Klebsiella* may exclude high risk-associated *Escherichia*. A) TaxUMAP visualization of samples followed by a BSI within the next week, no infection (grey), BSI by gamma proteobacteria (dark red), or enterococcal species (green); inset: TaxUMAP labeled by most abundant taxa as in figure 1I, histogram shows relative frequencies of blood isolate taxa. B) TaxUMAP of the mean a posteriori predicted BSI risk per sample from a Bayesian logistic regression comparing stool compositions between BSI cases and uninfected patients using the log_10_-relative ASV abundances of 10 predictor ASVs, the histogram shows the distribution of posterior risk predictions across all samples; inset: magnified region with the highest predicted risk samples. C) Receiver-operator characteristic of the Bayesian model. D) Posterior coefficient distributions (circle: mean, box: HDI50, whiskers HDI95). E) High risk samples are located in the Enterobacteriaceae dominated TaxUMAP region (top: TaxUMAP with samples labeled by the most prevalent taxon as in Figure 1I); ASV-3 and ASV-568 (increased risk) and ASV-36 (reduced risk) relative abundances across the TaxUMAP and in the magnified Enterobacteriaceae dominated region. Blood isolate organism abbreviations) EFVR: *E. faecium* vancomycin resistant, Esch: *Escherichia*, KP: *K. pneumoniae*, Ebct: *Enterobacter*, EF: *E. faecium*, P: *Pseudomonas*, SM: *Stenotrophomonas maltophilia*, EVR: *Enterococcus* vancomycin resistant, EFlis: *E. faecalis*, K: *Klebsiella*, C: *Citrobacter*; abbreviations in ROC, C) TPR: true positive rate, FPR: false positive rate.

**Figure 4.**
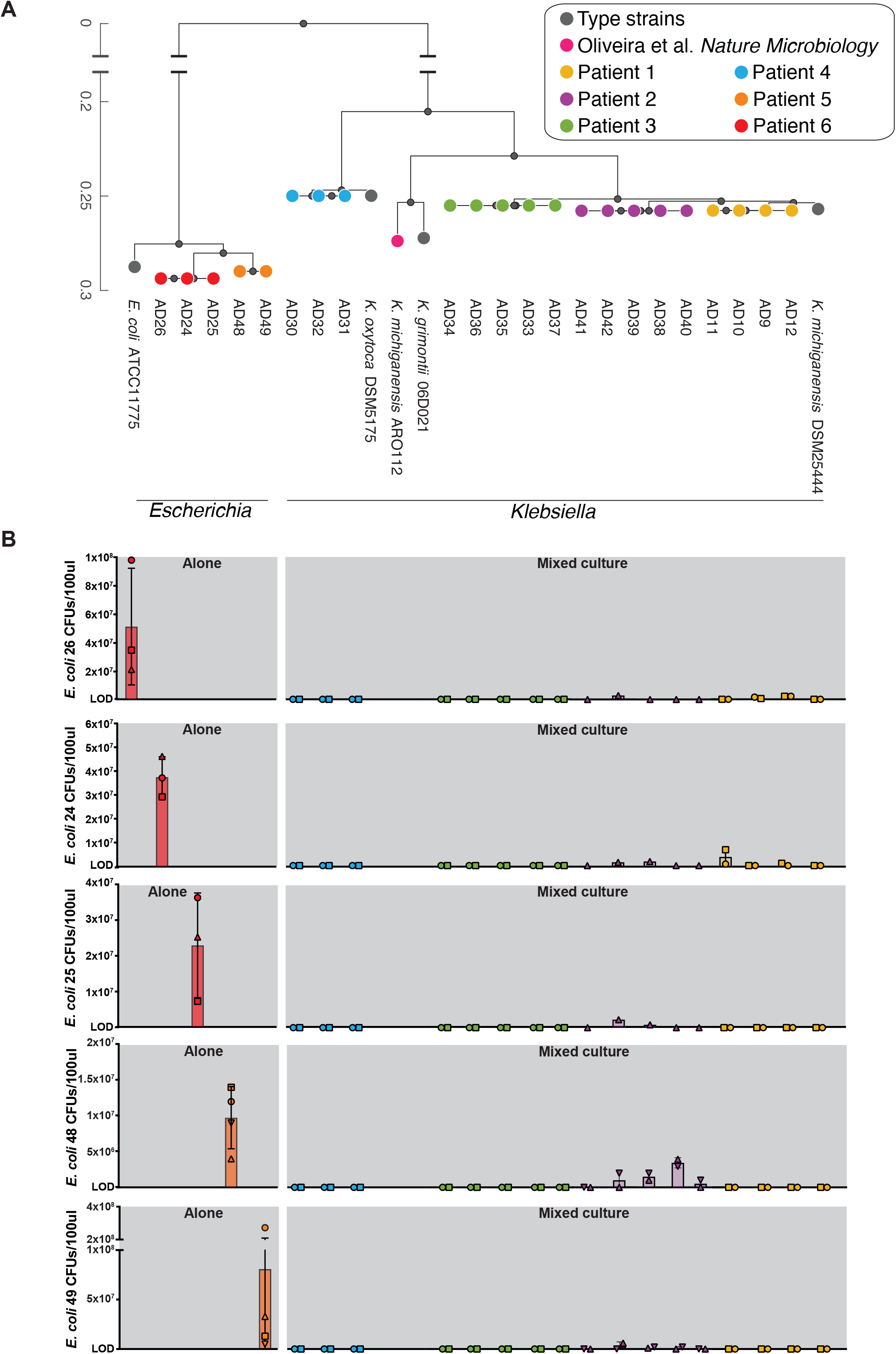
Ecological interference by ASV-36-*Klebsiella* isolates prevents expansion of BSI-associated ASV-3-*Escherichia* isolates in *in vitro* competitive exclusion assays. A) Phylogenetic tree representing 17 ASV-36-*Klebsiella* isolates and five ASV-3-*E. coli* isolates; colors indicate isolates coming from the same patient. See also **Figure S9** and **Table S1**. B) Results of ecological invasion assays. Bars represent mean CFU counts of ASV-3-*E. coli* isolates after 12h of incubation. Rows correspond to each of the ASV-3-*E. coli* isolates: red and orange bars represent isolates grown in absence of a resident ASV-36-*Klebsiella* community; other bars correspond to ASV-3-*E. coli* CFUs grown in presence of different ASV-36-*Klebsiella* isolates. The colors of the bars and their order indicate which of the 17 ASV-36-*Klebsiella* strains represented on the tree in A) was used as a competitor in the assay. Error bars: standard deviation, symbols indicate three independent experiments. CFU: colony forming unit.

Interestingly, the model revealed that ASV-36 was negatively associated with infection (**Figure 3C**). Visual inspection of the TaxUMAP confirmed that ASV-36 is enriched in a low risk sub-region of the Enterobacteriaceae-enriched region (**Figure 3E**). ASV-3 and ASV-568, two Enterobacteriaceae taxa positively associated with infection, were highly abundant in directly adjacent regions. ASV-36 had made it to the list of 10 candidates because it was highly abundant preceding three BSI cases where the bacteria isolated from the patient’s blood were annotated as “*Klebsiella sp*.”. The 16S rRNA gene sequence of ASV-36 precisely matches several species of the *Klebsiella* genus, including *K. michiganensis* and *K. oxytoca* (**Figure S9**), but not *K. pneumoniae*.

### Experiments validate ASV-36-*Klebsiella* ability to exclude bacteria associated with high infection risk

Our analysis so far shows that ASV-36-*Klebsiella* contribute negatively to the infection risk model, and the TaxUMAP suggested this ASV may exclude other bacteria associated with higher risk of infection like *E. coli* (ASV-3) and *K. pneumoniae* (ASV-30). This finding was especially intriguing because two recent studies showed in mice that similar strains of *Klebsiella* could protect mice against pathogenic Enterobacteriaceae by mechanisms of nutrient competition (Oliveira et al., 2020; Osbelt et al., 2021). Our results together suggest ASV-36-*Klebsiella* could be an ecological competitor of other bacteria associated with high risk of infection in humans.

We sought to characterize ASV-36-*Klebsiella* beyond the resolution of 16S amplicon sequencing. We first analyzed the subset of 16 shotgun-sequenced samples from 4 unique patients containing ASV-36-*Klebsiella* with relative abundance ⩾1% and identified eight metagenomically-assembled genomes (MAGs, **Figure S9**). Two of those MAGs came from one patient and matched *K. michiganensis;* six other MAGs came from another patient and matched *K. grimontii*. We then obtained 17 strain isolates by plating on selective MacConkey agar stool samples containing high levels of ASV-36 originating from four different patients (**Table S1**). Whole genome sequencing allowed construction of a phylogenetic tree of the 17 ASV-36 isolates that additionally comprises type strains and the murine *K. michiganensis* ARO112 strain (Oliveira et al., 2020), as well as the high-risk associated ASV-3 isolates (**Figure 4A**). The tree shows that isolates from the same patient are clonal, indicating a larger inter- than intra-individual variability among strains subsumed by ASV-36-*Klebsiella*. A larger phylogenetic tree constructed from our isolates alongside all published genomes containing the ASV-36-*Klebsiella* 16S rRNA gene sequence shows that isolates from three patients colocalized in the branch defined mostly by *K. michiganensis* genomes, while isolates obtained from one patient positioned on the branch dominated by *K. oxytoca* genomes (**Figure S9**). These results indicate that ASV-36-*Klebsiella* represents several related species of *Klebsiella*, including *K. oxytoca* and *K. michiganensis*, but not *K. pneumoniae*.

The TaxUMAP atlas (**Figure 3E**) suggested a competitive exclusion between ASV-36-*Klebsiella* and other, potentially more virulent Enterobacteriaceae, such as ASV-3. If this was the case, therefore, an established community of ASV-36-*Klebsiella* would interfere with ASV-3 colonization. To validate this mechanism experimentally we isolated 5 different ASV-3 strains from two different patients’ stool samples (**Table S1**). Whole genome sequencing of each isolate confirmed them as strains of *E. coli*. To test if an established ASV-36-*Klebsiella* community could prevent ASV-3-*E. coli* colonization, as suggested by the TaxUMAP, we performed a simple experiment to test the minimum requirement for such competition. We first established communities of each of the 17 isolated strains *in vitro* by growing them in LB media to stationary phase. We then introduced ASV-3-*E. coli* and measured their growth compared to their growth in a sterile LB flask. If ASV-3-*E. coli* and ASV-36-*Klebsiella* shared an ecological niche, including similar or identical nutrient requirements, we would expect a growth reduction of ASV-3-*E. coli*. Indeed, after 12 more hours of incubation, ASV-3-*E. coli* reached densities of up to 10^8^ CFUs when unimpeded by an established ASV-36-*Klebsiella* (**Figure 4B**). But the counts dropped precipitously if their niche was previously colonized by ASV-36-*Klebsiella* (**Figure 4B)**. Out of the 17 ASV-36-*Klebsiella* isolates, 11 entirely prevented expansion of all five tested ASV-3-*E. coli* isolates (limit of detection: 1×10^6^ CFUs/100μl, see Methods). The 6 remaining ASV-36-*Klebsiella* isolates also strongly inhibited expansion of ASV-3-*E. coli*. In summary, every one of the tested ASV-36-*Klebsiella* isolates prevented expansion of ASV-3-*E. coli*. This, together with the findings from ARO112 in mice (Oliveira et al., 2020), suggests that the phenotype of ecological exclusion is conserved across a taxonomic group that includes *K. oxytoca* and *K. michiganensis*, potentially explaining the pattern seen on the TaxUMAP which suggests mutual exclusivity (**Figure 3E**). Therefore, an established low BSI-risk associated ASV-36-*Klebsiella* community may protect patients from the invasion by high BSI risk associated ASV-3-*E. coli*.

### *In vitro* competition reveals carbon sources that provide an advantage to ASV-36-*Klebsiella* over ASV-3-*E. coli*

In order to determine the conditions that would favor ASV-36-*Klebsiella* versus ASV-3-*E. coli*, we quantified their fitness ratio through direct competition in an array of nutrients *in vitro*. We competed the two strains in 95 different carbon sources (using BIOLOG Phenotype MicroArray-PM1 plates), which we inoculated with a mix of ASV-36 and ASV-3 at a starting ratio of 1:1, then we incubated them in both aerobic (common room conditions) and anaerobic conditions (inside an anaerobic chamber). The post-competition ratio measured after 24h (**Figure 5A-B**) showed that the majority of the tested carbon sources favored ASV-3 both in anaerobiosis and aerobiosis. However, the experiment revealed that the outcome of the competition depends on the environment: interestingly, in anaerobiosis ASV-36 had the advantage in 12 different carbon sources (**Figure 5A, Table S2**) but that number rose to 31 (**Figure 5B, Table S3**) in aerobic conditions (p<0.05 after multiple hypothesis correction, linear mixed-effects model). Ten of those carbon sources favored ASV-36 regardless of the presence of oxygen (**Figure 5A-B**, shown in blue), and this includes metabolites such as the sugars sucrose and maltose, known to be available to gut bacteria and important for their fitness (Jones et al., 2008; Townsend et al., 2019). To test if common chemical characteristics between carbon sources favor ASV-36 we correlated the competition index (calculated as ASV-36 to ASV-3 ratio) with the molecular weight and with the number of carbons for each carbon source; but these simple chemical features were not correlated with fitness differences (**Figure S10**). Interestingly, *in vitro* experiments showed that 11 of the carbon sources that favored ASV-3 in anaerobic conditions, changed to favoring ASV-36 in aerobic conditions (**Figure 5A-B**, shown in red). No carbon sources worked in the opposite direction, i.e. providing an advantage to ASV-36 in anaerobic conditions yet favoring ASV-3 in aerobic conditions. This suggested that the switch from anaerobic to aerobic environments provided an exclusive advantage to ASV-36. All together, these results suggest that perturbations which increase oxygen availability could increase the range of environmental conditions favoring ASV-36-*Klebsiella*.

**Figure 5.**
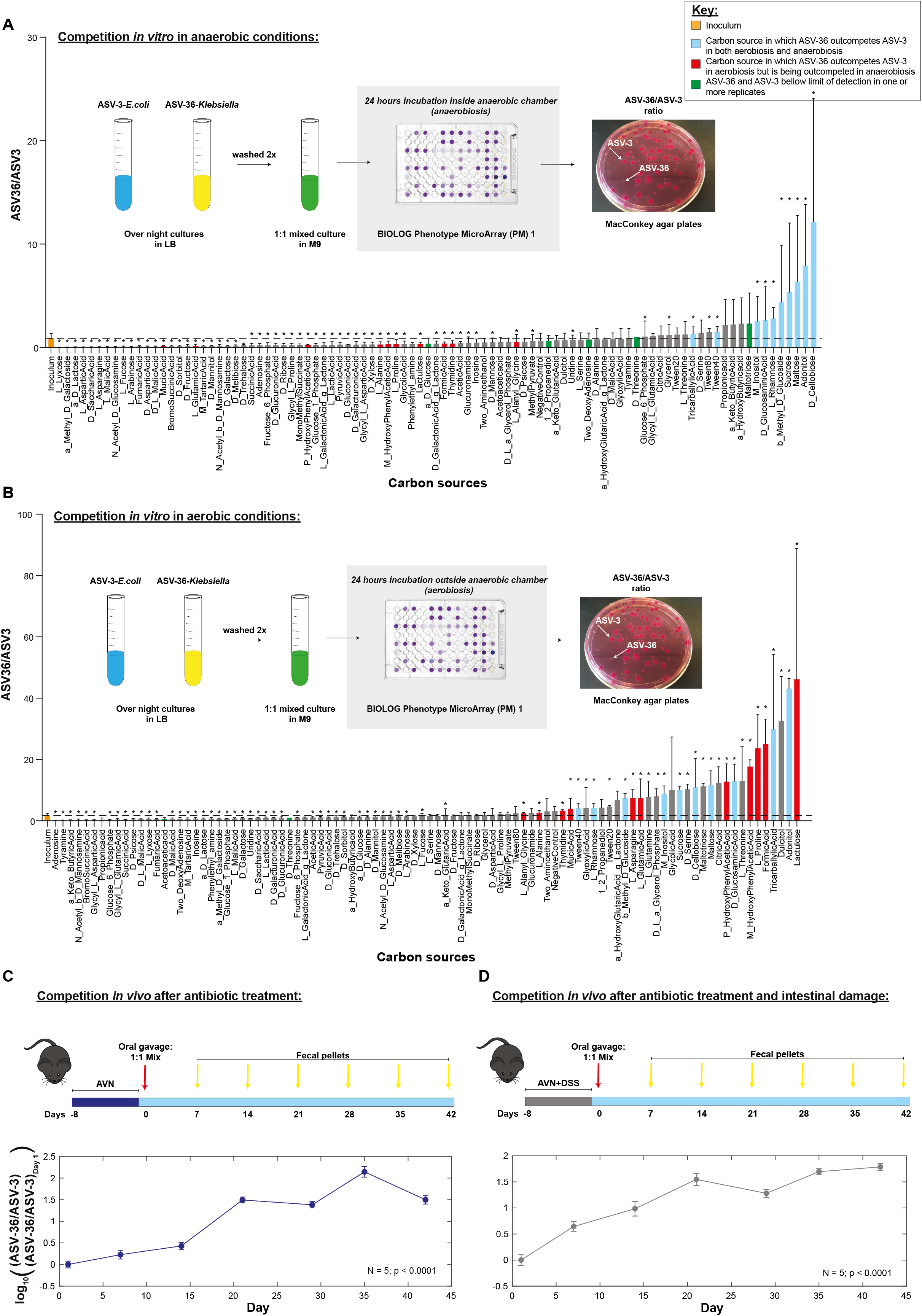
ASV-36-*Klebsiella* and ASV-3-*E. coli in vitro* competition results depend on the available nutrients and oxygen concentration, however conditions in the gut of antibiotic and DSS treated animals favor ASV-36. A) ASV-36-*Klebsiella* outcompetes ASV-3-*E. coli* in 12 different carbon sources in the absence of oxygen. Plot represents scheme of *in vitro* experiment and its results. Pure ASV-3 and ASV-36 overnight cultures were centrifuged and washed 2 times in M9 minimal media lacking carbon source, after which they were re-suspended and mixed in ∼1:1 ratio. The mixed culture in M9 media was used to inoculate BIOLOG PM1 plates that were then incubated inside of an anaerobic chamber. After 24h every well was plated on MacConkey agar plates, ASV-3 and ASV-36 colonies were counted and ASV-36 to ASV-3 ratio was determined. Although the majority of tested nutrients provided advantage to ASV-3, there are 12 different carbon sources in which ASV-36 can outcompete ASV-3. See also **Figure S10** and **Table S2**. Asterisk mark carbon sources in which ASV-36/ASV-3 ratio was significantly different from the ratio in the inoculum (p<0.05 after multiple hypothesis testing; linear mixed-effects model; N=3 independent experiments). B) ASV-36-*Klebsiella* outcompetes ASV-3-*E. coli* in 31 different carbon sources in the presence of oxygen. The performed experiment is similar to the one described in 5A, with the exception that the BIOLOG PM1 plates were incubated in aerobic conditions, which led to a higher number (31) of carbon sources that provided advantage to ASV-36 as compared to anaerobic conditions. While some carbon sources (marked blue), provided advantage to ASV-36 no matter the condition, some others (marked red) provided the advantage in aerobic conditions but in anaerobiosis lead to ASV-36 being outcompeted by ASV-3. ASV-36 and ASV-3 were both under limit of detection in one or more replicates for carbon sources marked in green. See also **Figure S10** and **Table S3**. Asterisks mark carbon sources in which the ASV-36/ASV-3 ratio is significantly different from the ratio in the inoculum (p<0.05 after multiple hypothesis testing; linear mixed-effects model; N=3 independent experiments). C) ASV-36-*Klebsiella* outcompetes ASV-3-*E. coli in vivo* in a mouse model of antibiotic-induced co-colonization. Plot contains a schematic representation of performed experiment and its results. After antibiotic cocktail treatment (ampicillin, vancomycin and neomycin-AVN) mice were orally gavaged with 10^7^ CFUs of 1:1 mix of both strains. In co-colonized mice ASV-36-*Klebsiella* showed competitive advantage over ASV-3-*E. coli* which was reflected by the significant increase in ASV-36/ASV-3 ratio during the course of the experiment (p<0.0001; N=5 mice; linear mixed-effects model). D) Dextran sodium sulfate (DSS) treatment leads to a similar advantage of ASV-36-*Klebsiella* over ASV-3-*E. coli in vivo*. Another group of animals was treated with the same antibiotic cocktail and dextran sodium sulfate (AVN+DSS; schematic representation). Additional treatment with DSS resulted in similar increase in the ratio of ASV-36 to ASV-3 with respect to the first day post-infection (p<0.0001; N=5; linear mixed-effects model): There were no significant differences in rates of change between the group of mice that received AVN alone as compared to mice that received AVN+DSS suggesting that further damage of the intestinal epithelium provoked by DSS does not affect the result of the competition.

### ASV-36-*Klebsiella* are more competitive *in vivo*

Our *in vitro* assays showed that environmental conditions could alter the outcome of competition between ASV-36 and ASV-3. The implications of this context-dependent fitness in patients’ microbiomes are difficult to extrapolate: nutrients available in the gut depend on several factors including diet (David et al., 2014; Johnson et al., 2019) and the microbiome composition of other gut bacteria that will also consume and release metabolites (Rakoff-Nahoum et al., 2016; Ze et al., 2012). Anaerobiosis itself can be impacted by several factors, including the host’s intestinal inflammation and epithelial damage (Chanin et al., 2020; Litvak et al., 2018; Rigottier-Gois, 2013), which is frequent in allo-HCT patients, and could raise the concentration of oxygen in the lumen.

To assess the competition *in vivo*, we carried out experiments in mice (**Figure 5C**). We treated the mice with a cocktail of antibiotics (ampicillin, vancomycin and neomycin: AVN), and then inoculated them with either mono-cultures of ASV-3-*E*.*coli*, ASV-36-*Klebsiella* or a mixed culture of both. Mono-colonized mice showed that both ASV-3 and ASV-36 could colonize the gut after AVN treatment to achieve robust and stable levels of colonization. The mice colonized with the mixture, however, showed that ASV-36-*Klebsiella* had a competitive advantage (**Figure 5C**). The ratio of ASV-36 to ASV-3 grew steadily during the 42 day experiment, with a fitness advantage of *f*=0.13/day (±0.06, p=0.00002). To determine if the advantage of ASV-36 is impacted by host-induced environmental perturbations, we conducted an additional experiment (**Figure 5D**) where we treated the mice with antibiotics and dextran sodium sulfate (DSS). DSS is a model for inflammatory bowel disease (IBD) that leads to increased oxygenation of the gut, perturbs the intestinal barrier function and increases intestinal permeability more so than what is typically observed in IBD patients (Cochran et al., 2020); DSS treatment thus constitutes a major perturbation to epithelial homeostasis that is also expected in chemotherapeutically treated cancer patients. In AVN+DSS treated mice, we observed a slight increase in the relative fitness of ASV-36 to *f*=0.14 (**Figure 5D**). Taken together, these results show that ASV-36-*Klebsiella* outcompetes ASV-3-*E. coli in vivo*, when inoculated simultaneously, and that the competitive advantage is robust to extreme environmental perturbations.

## DISCUSSION

Here we present TaxUMAP, a taxonomy-aware tool to display large longitudinal clinical microbiome data and identify causal microbiome effects on human health. We used TaxUMAP to produce an atlas from data of patients hospitalized to receive allo-HCT at Memorial Sloan Kettering Cancer Center (Liao et al., 2021; Yan et al., 2022). Hospitalized transplant patients such as these provide a unique opportunity for new ecological insight for four main reasons: First, their gut microbiota can undergo extreme perturbations that span much wider compositional states than healthy individuals. Second, hospitalized patients are closely monitored, and the rich metadata enables us to functionally interpret the compositional states. Third, high incidences of microbiome related, health relevant events such as BSIs empower statistical analyses, here used to establish data-driven definitions of microbiome homeostasis and dysbiosis. Fourth, HCT are conducted in a highly scheduled fashion with government mandated clinical data collection offering an opportunity for causal inference similar to “natural experiments”.

We saw that across all our samples, antibiotic-exposed and not exposed, the most diverse gut microbiota samples—which tended to be those collected prior to antibiotic treatment started—also harbor the most diverse repertoire of antimicrobial resistance genes. This interesting observation may result from fierce competition between microbes that wield antibiotics to kill other species and have counteracting resistance mechanisms as part of the microbial warfare arsenal (Donia et al., 2014; Mavridou et al., 2018). We saw that exposure to antibiotics was associated with a reduction in both microbiota diversity and the number of antimicrobial resistance genes. We hypothesize that the correlation between the number of resistance genes and microbiota diversity results from the strong selection for specific resistances that dominate the population due to antibiotic treatment. Our results have therefore important implications for the interpretation of metagenomic counting of antimicrobial resistance genes in the gut microbiome of hospitalized patients such as our cohort: we demonstrate that antibiotics select for specific genes rather than expanding the antibiotic resistance gene repertoire in the microbiome (Montassier et al., 2021).

We also saw that diversity correlates with total bacterial load. Diversity and load are both depleted to extreme degrees in allo-HCT patients. But we only saw a positive correlation between the two when we corrected for stool consistency. Among liquid stool samples, for example when patients had diarrhea, diversity is anti-correlated with total bacterial load. This observation confirms that stool consistency is an important—though often neglected— microbiome variable (Vandeputte et al., 2016).

Intestinal domination is a risk factor for bloodstream infection in immunocompromised cancer patients receiving allo-HCT (Stoma et al., 2021; Taur et al., 2012; Ubeda et al., 2010; Zhai et al., 2020). The TaxUMAP atlas confirms this finding visually. The Bayesian risk model that we developed to predict infection risk from a microbiota composition revealed clear differences between different gram negative enterobacteria. This model led us to find the ASV-36 associated with lower risk of infection. We used the atlas to select stool from our sample bank and isolate representative strains. We genotyped 17 ASV-36 isolates from patients’ samples, alongside isolates for high-risk associated enterobacterial sequence variants. ASV-36-*Klebsiella* comprises several species, including *K. michiganensis* and *K. oxytoca*. Importantly, all sequenced ASV-36 isolates were clearly distinct from the genome of *Klebsiella pneumoniae*, a known human pathogen which in our data is ASV-30.

Related bacteria can form nutrient-exchanging food webs with potentially mutually beneficial effects on their growth (Rakoff-Nahoum et al., 2016); however, most microbiome species and bacterial taxa in general are predicted to compete with one another (Bucci et al., 2012; Coyte et al., 2015; Foster and Bell, 2012; Foster et al., 2017; Palmer and Foster, 2022). Such facilitation or interference could reveal tangible microbiome engineering principles if the direction of interactions could be identified (Palmer and Foster, 2022). We investigated facilitation or interference between strains with a plausible causal health effect on hospitalized patients. The TaxUMAP atlas had suggested that ASV-36-*Klebsiella*, which were associated with a reduced risk for BSIs, appeared to exclude high BSI-risk associated ASV-3-*E. coli*. We thus tested 85 unique combinations of strains to establish if high-risk associated ASV-3 isolates would establish themselves in an existing community of ASV-36-*Klebsiella* and found that the exclusion expected from the TaxUMAP visualization could be confirmed experimentally.

Such an exclusion effect might be expected when cell densities in an established culture are already high, and many nutrients might have been consumed such that newly arriving stains fail to grow on this spent medium; but only when the competing strains share nutrient requirements and are able to utilize the same available nutrients rather than facilitate each other by cross feeding (Rakoff-Nahoum et al., 2016). In fact, when we tested the growth capacity of ASV-36-*Klebsiella* and ASV-3-*E*.*coli* in different nutrient sources (BIOLOG PM1 plate) we saw that there is a series of carbon sources in which both strains can grow. While the majority of the tested carbon sources provided advantage to ASV-3, some boosted the growth of ASV-36, and others had an effect that was dependent on oxygen presence. Among carbon sources that boosted ASV-36 growth in both aerobic and anaerobic conditions are beta-Methyl-D-Glucoside, D-cellobiose, sucrose, inositol, that were previously described to provide growth advantage to protective *K. oxytoca* strains over multidrug-resistant *K. pneumoniae* (Osbelt et al., 2021). Interestingly, *in vitro* experiments suggest that increase in aerobiosis results in ASV-36 outcompeting ASV-3 in more carbon sources than what was observed in anaerobic conditions. This might be relevant for our patient cohort that undergoes extensive antibiotic treatment and often suffers gut inflammation, both of which were shown to increase the availability of the oxygen in the gut of mice (Cevallos et al., 2021; Rivera-Chávez et al., 2016), and which ultimately leads to the bloom of bacteria from the family Enterobacteriaceae. Thus, our *in vitro* results demonstrate that ASV-36 and ASV-3 might compete for a niche and that this competition might be tilted in favor of ASV-36 in a more oxygenated environment such is presumably the gut of our patients. To test whether the niche competition would occur *in vivo*, we performed co-colonization experiments in antibiotic treated mice. Our results show that ASV-36 outcompetes ASV-3 *in vivo*. We also wanted to see if epithelial damage, which often happens in our patient cohort, could alter the result of the competition. However, it seems that in the animals that have been receiving DSS, which is known for provoking epithelial damage, ASV-36 have the same advantage over ASV-3 as what was observed in mice treated with antibiotics exclusively, suggesting that even in the case of this extreme perturbation of the gut ASV-36-*Klebsiella* outcompetes ASV-3*-E. coli*. In summary, despite the results from *in vitro* experiments, where it seemed like ASV-3 might be fitter than ASV-36, *in vivo* experiments demonstrate that ASV-36 outcompetes ASV-3 both in antibiotic treated animals, and those with damaged epithelial barrier. Our results agree with recent mouse studies where related strains of *Klebsiella* could competitively exclude pathogenic Enterobacteriaceae (Oliveira et al., 2020; Osbelt et al., 2021). Our study supports that the same mechanism occurs in patients: presence of a low-risk gram negative enterobacterium could exclude subsequent establishment of pathobiont enterobacteria in a perturbed microbiome and thereby reduce risk of infection.

Taken together, the TaxUMAP atlas provides a new way to mine a longitudinal clinical microbiome dataset for causal microbiome effects on human health. The tools published here can continue to be applied to the vast data of allo-HCT microbiome states that we generated at MSKCC (Liao et al., 2021; Yan et al., 2022), and address other research questions besides the ones proposed here. The same method can also be applied to other longitudinal microbiome datasets, and generate new hypotheses on causal microbiome effects on human health.

## ACKNOWLEDGEMENTS

This work was supported by the National Institutes of Health (NIH) grants DP2 AI164318-01 to J.S.; U01 AI124275 to E.P. and J.B.X.; R01 AI137269 to J.B.X. and Y.T.; NHLBI NIH Award K08HL143189, the MSKCC Cancer Center Core Grant NCI P30 CA008748, the Parker Institute for Cancer Immunotherapy at Memorial Sloan Kettering Cancer Center to J.U.P.; NIH R01 AI14862303, NIH R01 AI143757, CDC BAA 75D30118C02921, Sloan Foundation Fellowship, V Foundation Fellowship, Stand Up 2 Cancer award 6206 to A.S.B.; National Cancer Institute award numbers R01-CA228358, R01-CA228308, P30 CA008748 MSK Cancer Center Support Grant/Core Grant and P01-CA023766, National Heart, Lung, and Blood Institute (NHLBI) award number R01-HL123340 and R01-HL147584, National Institute of Aging award number P01-AG052359, Starr Cancer Consortium, Tri Institutional Stem Cell Initiative, The Lymphoma Foundation, The Susan and Peter Solomon Divisional Genomics Program, Cycle for Survival, and the Parker Institute for Cancer Immunotherapy to M.R.M.v.d.B.

## AUTHOR CONTRIBUTIONS

J.S. and J.B.X. conceived of the study. A.D. obtained fecal isolates, sequenced their genomes and validated the findings experimentally *in vitro* and *in vivo*. J.S., B.P.T. conceived of and J.S., G.A.H., C.D. implemented the TaxUMAP algorithm. J.S. implemented the Bayesian analyses. B.P.T. implemented 16S data processing pipelines. C.L. performed the metagenome assemblies with the help of the lab of A.S.B, J.Y. performed the genomic comparisons of isolates. J.S. produced figures 1-3 and A.D. figures 4 and 5. J.U.P. and Y.T. helped with the provision and interpretation of clinical metadata. All co-authors read and contributed to the manuscript.

## METHODS

### Resource availability

#### Lead contact

Further information and requests for resources and reagents should be directed to and will be fulfilled by the lead contact, Dr. Joao B. Xavier.

### Materials availability

All unique/stable reagents generated in this study are available from the Lead Contact without restriction.

### Data and code availability

The TaxUMAP algorithm is available on GitHub (https://github.com/jsevo/taxumap). Sequencing files generated by whole genome sequencing of bacterial isolates have been submitted to NCBI and are publicly available (**Table S1**).

### Isolation and sequencing of ASV-36-*Klebsiella and* ASV-3-*E. coli*

For isolation, patients’ samples with high relative abundance (>90%) of desired ASVs were plated on MacConkey agar plates (BD MacConkey II Agar). Plates were incubated in aerobic conditions at 37ºC for 24-48h. Individual colonies were picked, and Sanger sequenced to confirm their identity based on 16S rRNA gene. For whole genome sequencing (WGS) a single colony was grown overnight in LB liquid media. Next day 1ml of the culture was centrifuged at 15 000rpm for 5 minutes and pellet was subjected to DNA extraction with Qiagen Fast DNA Stool Mini Kit. Nextera XT DNA Library Preparation Kit was used for library generation according to the manufacturer’s instructions. Libraries were quantified with Quant-iT™ dsDNA Assay Kit, normalized and sequenced using MiSeq Reagent Kit V3. Comprehensive Genome Analysis tool in PATRIC (Wattam et al., 2017) was used for sequence processing: genomes were assembled using SPAdes (Bankevich et al., 2012) and annotated using RAST Tool Kit (Brettin et al., 2015). Phylogenetic Tree tool in PATRIC was used for codon tree generation from 1,000 randomly selected single-copy genes by using the RAxML (Stamatakis, 2006). Genome sequences are available at NCBI (**Table S1**).

### Phylogenetic tree of ASV-36-*Klebsiella*

The *Klebsiella* genomes that contain the same 16S gene sequence as ASV-36 were obtained from NCBI. The previously studied strains *K. oxytoca* CAV1374, *K. oxytoca* KCTC1686, *K. michiganensis* strain ARO112, and *K. sp*. Kd70 TUC-EEAOC were also included and they belong to ASV-36 as well. All *Klebsiella* genomes including the metagenome-assembled genomes (MAGs) and our isolated ASV-36-*Klebsiella* strains were annotated using prokka (Seemann, 2014) and the core genome alignment was generated by roary (Page et al., 2015) to further construct the phylogenetic tree using FastTree2 (Price et al., 2010). The *K. pneumoniae* reference genome *K. pneumoniae subsp. pneumoniae* HS11286 that does not belong to ASV-36 was used to root the tree. The species information was taken from the NCBI genome annotation.

### Assembly of *Klebsiella* genomes from Shotgun samples

The draft genomes were assembled from 16 shotgun samples (1042X, FMT.0009X, FMT.0009Y, FMT.0103Y, 668DD, 668T, 668EE, 668Y, 668CC, 668FF, 668X, 668GG, 668Z, 668W, 668MM, 668BB; accessible from Bioproject PRJNA545312) which contain at least 1% of ASV-36-*Klebsiella* based on 16S amplicon sequencing. The assembly pipeline was downloaded from the Bhatt lab github repository (https://github.com/bhattlab/bhattlab_workflows) and installed locally. Briefly, the computational workflows used Megahit (Li et al., 2015) for genome assembly and both Metabat2 (Kang et al., 2019) and CONCOCT (Alneberg et al., 2014) for metagenomic binning.

### Ecological invasion experiments

Competitive exclusion of ASV-3-*E. coli* isolate invasion by resident ASV-36-*Klebsiella* was tested by challenging established ASV-36-*Klebsiella* communities with ASV-3-*E. coli*. To generate an established ASV-36-*Klebsiella* community to be invaded by ASV-3-*E. coli*, ASV-36 strains were grown in liquid LB medium from fresh overnight culture until reaching stationary phase. To do this ASV-36-*Klebsiella* strains 9, 10, 11, 12, 30, 31, 32, 33, 34, 35, 36, 37 were grown for 12h and strains 38, 39, 40, 41, 42 were grown for 20h. The ASV-36-*Klebsiella* incubation times were selected based on the different growth rates among our isolates that resulted in some of the isolates reaching the stationary phase earlier than the others. Once the ASV-36-*Klebsiella* isolates reached the stationary phase they were inoculated (1:50) with ASV-3-*E. coli* isolates in the exponential phase of growth. As a control, LB medium not containing ASV-36-*Klebsiella* culture was also inoculated with ASV-3-*E. coli*. After 12h of incubation in aerobic conditions at 37ºC and with shaking cultures were plated on MacConkey agar and colony forming units (CFUs) of ASV-3-*E. coli* in mixed cultures were compared to those observed when ASV-3-*E. coli* were grown alone. All ASV-36-*Klebsiella* isolates were tested against all ASV-3-*E. coli* isolates making a total of 85 tested combinations. The limit of detection was 1×10^6^ CFUs/100μl. This limit was due to the fact that detection of both competing strains was done on the same agar plate. Overgrowth of ASV-36-*Klebsiella* on more concentrated plates prevented a higher resolution that could allow a lower limit of detection of ASV-3-*E. coli*.

### Ecological competition in different nutrient environments (BIOLOG assays)

To test in which conditions ASV-36-*Klebsiella* might outcompete ASV-3-*E*.*coli* we setup *in vitro* competition assay that tested their growth in different carbon sources in BIOLOG Phenotype MicroArray (PM) 1 in presence or absence of oxygen. A single colony of ASV-36 (AD9 strain) and ASV-3 (AD24 strain) was used to inoculate 5ml of LB, after which cultures were grown overnight at 37ºC and with shaking. Next day, both cultures were centrifuged at maximum speed for 5 minutes and re-suspended in M9 minimal media lacking carbon source. This washing step was performed twice. After the third and final centrifugation step, strains were re-suspended in M9 minimal media until reaching the same OD600 absorbance of 0.05. ASV-36 and ASV-3 were mixed in 1:1 ratio and 100ul was aliquoted in each well of the BIOLOG PM1 plate. BIOLOG PM1 plate contains 95 different carbon sources and allows for assessing the growth capacity of tested strains on each one of them. Plates were incubated inside of plate readers at 37ºC and with shaking both inside the anaerobic chamber (anaerobiosis) and on the bench top (aerobiosis). After 24h aliquots from each well were serially diluted and plated on MacConkey agar plates. ASV-36 and ASV-3 colonies were counted and ASV-36/ASV-3 ratio was determined for each BIOLOG well. The initial inoculum was plated the same way to determine the starting ASV-36/ASV-3 ratio. Experiment was done in triplicate. The limit of detection for aerobic growth was 1×10^6^ CFUs/100μl, and for anaerobic growth 1×10^5^ CFUs/100μl. Generalized linear mixed-effects model implemented in Matlab 2021b was used to determine which carbon sources had ASV-36/ASV-3 ratio significantly different than the one detected in corresponding starting inoculum.

### *In vivo* competition assays

In order to test the outcome of ASV-36-*Klebsiella* and ASV-3-*E. coli* competition *in vivo* the following experiment was performed. We treated 6-8 weeks old C57BL/6J female mice with a cocktail of ampicillin (0.5g/l), vancomycin (0.5g/l) and neomycin (1g/l) for one week in drinking water. Antibiotics were changed once during the course of the treatment. Animals were single housed in autoclaved cages. Autoclaved water supplemented with antibiotics and 5053 irradiated food was provided *ad libitum*. The day after antibiotic cessation animals were orally gavaged with 10^7^ CFUs of ASV-36-*Klebsiella*, ASV-3-*E. coli* or 1:1 mix of both strains in PBS. The levels of colonization were monitored by collecting fecal pellets and plating them in MacConkey agar plates. To test whether profound epithelial damage could alter the result of competition, one group of animals was treated with the same antibiotic cocktail and 3% dextran sodium sulfate (DSS) in the drinking water for a week. The water with antibiotics and DSS was changed once, as previously mentioned. Animals were orally gavaged with 10^7^ CFUs of 1:1 mix of ASV-36 and ASV-3 the day after treatment withdrawal. The levels of colonization were monitored by collecting fecal pellets and plating them in MacConkey agar plates. Generalized linear mixed-effects model implemented in Matlab 2021b was used to determine the change in ASV-36/ASV-3 ratio over the course of the experiments.

### *vanA* PCR from rectal swabs

Data obtained from clinical records; rectal swabs are routinely collected and analyzed for presence of the *vanA* gene via PCR on the same day a corresponding stool sample was taken.

### Alpha-diversity calculation

We calculate the inverse Simpson index (*s*) in sample *i* based upon the *n* relative ASV abundances (*p*) using the following equation: 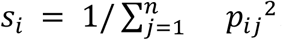.

### Pseudo time visualization of samples in clinical phases

While HCT is a strictly planned therapy, per patient variability arises. This leads to different clinical and medication exposures per patient per day, and for visualization purposes of trends over treatment regimens, we harmonize this by introducing a pseudo time that splits each patient’s treatment timeline into 9 segments corresponding to 3 points per each of the 3 clinical phases. We provide an antibiotic exposure summary table (**Table S4**) for each of the three phases, and visualize antibiotic exposure per treatment day as well as per pseudo timepoints (**Figure S11**).

### The TaxUMAP algorithm

The TaxUMAP algorithm is publicly available as a package for the Python programming language and as a command line application. We provide an extensive online tutorial on https://github.com/jsevo/taxumap, including data used in the manuscript, as well as smaller published data sets for exploration. The algorithm starts by calculating the sample-by-sample distances on ASV relative abundances. The algorithm allows users to choose from a range of different distance metrics; here, we used the cityblock distance *d* on relative abundances throughout, defined as 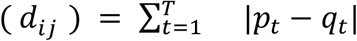 between samples *i* and *j* represented by the composition vectors *p* and *q, i*.*e*. the element-wise absolute difference in each taxon *t* out of all *T* taxa. Then, the algorithm calculates additional sample-by-sample distance metrics based on user-chosen taxonomic aggregations of the relative abundances of bacterial taxa. For example, the sample by sample distances at the genus level are calculated by first summing the relative abundances of all ASVs belonging to the same genus and then calculating the L1-norm (or user defined distance metric) across all samples. The ranges of the sample-by-sample cityblock distances are the same for each taxonomic aggregation, but their magnitudes are on average equal or smaller at higher taxonomic aggregations. The TaxUMAP algorithm allows users to scale different taxonomic aggregations by assigning a weight to each aggregation; here we chose to weight distances at ASV, family, and phylum level equally, discarding distances at other levels. We also set this as the default implementation and explain in the documentation and tutorials how to change these settings (Figure S12). For example, a researcher may be less interested in phylum-level differences between samples but choose to weigh more heavily how dissimilar samples are with respect to their genus composition. The choice of the default implementation was inspired by the traditional importance of phylum level difference in microbiome samples (Ley et al., 2006), the reported conservation of important traits at the family level (Goldford et al., 2018), and the ASV level as the highest resolution data available to us. Before applying the UMAP algorithm (McInnes et al., 2018b), the weighted distance matrices are added. The UMAP algorithm is then applied with a minimum distance of 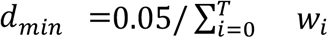 where *w*_*i*_ is the weight at taxon level *i* of *T* taxonomy levels, 120 neighbors, 10^3^ epochs. Each of these parameters can be altered by the user. We provide recommendations on the minimum distances and neighborhoods in the online documentation.

### Identification of risk-associated ASVs

To analyze the associations between the microbiome bacterial composition and identified bacteria of a BSI, we sought to associate relative ASV abundances with a risk score. For this we took the following steps: 1) match BSI organism to its bacterial family, 2) identify a list of candidate translocating ASVs in BSI-preceding stool samples, 3) analyze the risk of the ASVs to cause bacteremia by comparing their abundances in BSI cases vs. control samples.

#### Step 2) Identification of candidate taxa

We chose ASV candidates from among ASVs belonging to the same family as the BSI causing bacterium. We selected ASVs that were most abundant and/or increased the most in the seven days prior to a BSI event. The resulting list includes specific candidate translocator ASVs. We appended a single ASV, ASV_6997, to this list corresponding to a singleton *Citrobacter* BSI which could not be analyzed as the other, repeatedly observed BSIs. Manual inspection showed that the corresponding stool sample prior to the *Citrobacter* infection (sample 727C) was enriched for an Enterobacteriaceae ASV, ASV_6997, the sequence of which was mapped to *Citrobacter sedlakii* via BLAST.

#### Step 3) Matching case and control samples from uninfected patients

We created a matched case-control data set: case samples were the stool samples closest in time prior to a BSI event; controls were samples from patients without recorded BSI event chosen such that the sex of the patient and the clinical phase of the HCT therapy during which a sample was collected matched; we chose 4 controls for every case sample.

#### Step 3) Bayesian logistic regression

We performed a Bayesian logistic regression using the matched cases-control data with a constrained intercept, *a*, of -1.38, *i*.*e*. *inverse logit*(*a*) = 0.2, such that our model assumes the defined case-control probability *a priori*, with standard Normal predictor coefficient prior distributions. We implemented the model using the pymc3 package for the Python programming language and sampled 4 independent posterior chains using the No-U-turn sampling method (Homan and Gelman, 2014); the analysis code alongside the case-control data is provided in Supplementary file S1. For posterior predictions, we used the relative abundances of predictor taxa in a given sample and calculated the mean predicted probability of BSI from 10,000 posterior samples with an intercept correction *a*\* = *a*_*p*_, − *log*((1 − *τ*)/*τ* * ŷ/(1 − ŷ)), where *ŷ* is the fraction of cases, *a*_*p*_ the posterior intercept estimate, and *τ* the population incidence rate of BSIs (King et al., 2010).

## SUPPLEMENTAL INFORMATION TITLES AND LEGENDS

**Figure S1. Log-diversity of the bacterial microbiota is positively correlated with the summed relative abundances of obligate anaerobe taxa**. Related to **Figure 2G**. Blue: regression line with 95% CI from a linear model, slope: 0.92, p<10^−4^.

**Figure S2. The relationship between stool consistency and bacterial density and diversity**. Related to **Figure 2I**. A) Liquid and semi-formed stool samples have significantly lower bacterial densities than found in formed stool (p<10^−10^; two-sided t-tests). B) Formed and semi-formed stool samples show a positive relationship between diversity and total bacterial counts, whereas a negative relationship is found in liquid stool samples; 1,000 regression lines sampled from the joint posterior for formed (orange), semi-formed (green) and liquid (blue) stool samples, and corresponding posterior regression line means (black lines, beta: slope values) from a Bayesian mixed effects model with random intercepts and slopes per consistency category.

**Figure S3. Start locations of volatile switches (volatility >= 0.9) are found across the TaxUMAP**. Related to **Figure 2J**.

**Figure S4. Dominated states persist only partially due to antibiotic treatment pressure**. Related to **Figure 2J**. A) Percentage of patients that received their last antibiotic on the specific day relative to the day of transplant indicated on the x-axis; N=1,172 patients. B) Correlation between microbiome diversity (log_10_ of the inverse Simpson index) and days that passed since the last antibiotic administration reveals the average diversity doubling time of 44 days. Red line: slope and intercept from linear mixed-effects model. (N=853 samples, p<0.05, linear mixed-effects model) C) Half-life for each of the eight most prevalent dominating states varies highly. The plot on the left represents the percentage of cases with one of the top eight domination states. The plot on the right represents domination half-life calculated by a linear mixed-effects model, and its lower bounds. IS: inverse Simpson index; ASV: amplicon sequence variant.

**Figure S5. More diverse samples harbor more unique antimicrobial resistance phenotypes (blue: regression line from a linear model with 95% CI, p<10**^**-5**^). Related to **Figure 2K**.

**Figure S6. The effect of antibiotic treatments on microbiome diversity and presence of ARGs**. Related to **Figure 2F** and **2K**. A) The association coefficients between antibiotic treatments on bacterial diversity (log_10_ of the inverse Simpson index) as estimated by a linear mixed-effects model; circles: maximum likelihood estimate, error bars: 95% CI. The analysis revealed four antibiotics as significantly associated with the loss of bacterial diversity (marked in red intravenous piperacillin/tazobactam, intravenous meropenem, oral vancomycin and oral metronidazole). (Sample size N for each treatment stated in parenthesis next to the name of the antibiotic, p<0.05, linear mixed-effects model). B) Administration of these four antibiotics associated with the loss of bacterial diversity leads to an increase in a subset of the total range of ARGs. X axis represents ARG gene index sorted by the association strength and y axis represents Spearman correlation coefficient between ARG and diversity. Marked in yellow are the ARG genes whose number rises upon administration of each of the four antibiotics. (N=640 ARG genes, p<0.05 after multiple hypothesis testing, Spearman correlation). IS: inverse Simpson index; ARG: antibiotic resistance gene; CI: confidence interval.

**Figure S7. *vanA* gene detected in our patient cohort matches the *vanA* gene from *E. faecium***. Related to **Figure 2L**.

**Figure S8. The relationship between bacterial diversity and presence of *vanA* gene in rectal swabs**. Related to **Figure 2L**. A) Diversity in samples corresponding to positive *vanA* detection in PCRs from rectal swabs is significantly lower (p<10^−5^, Wilcoxon rank sum test, first sample per *vanA* status and per patient if multiple samples available, *vanA* negative: n=508, *vanA* positive: n=195). B) Positive *vanA* rectal swabs during phase I predict more severe loss of diversity in phases II and III, and higher *Enterococcus* abundances in phases II and III; if multiple samples per patient and clinical phase were available, the minimum observed diversity and maximum observed *Enterococcus* abundances were chosen in patients of either *vanA* group. P values from Wilcoxon rank sum tests; *vanA* negative in phase I: n=644, *vanA* positive in Phase I: N=85.

**Figure S9. Phylogenetic tree representing the relationship between ASV-36-*Klebsiella* isolates and other publicly available *Klebsiella* genomes**. Related to **Figure 4**. The tree was built using genomes deposited at NCBI database whose 16S rRNA gene matched the 16S sequence of ASV-36-*Klebsiella* exactly, 8 MAGs constructed from our patients’ samples and the genomes of ASV-36 closest neighbors (*i*.*e. K. oxytoca* CAV1374 and *K. oxytoca* KCTC1686) and the genome of *K. michiganensis* ARO112 described in a previous study (Oliveira et al., 2020). The tree was rooted with a genome of the pathobiont *K. pneumoniae subsp. pneumoniae* HS1128 strain. Colors correspond to taxonomic classification at the level of species.

**Figure S10. Correlation between ASV-36 to ASV-3 ratio and molecular weight and carbon numbers of each carbon source in BIOLOG PM1 plate**. Related to **Figure 5A** and **5B**.

**Figure S11. Antibiotic administration exposures**, over time. (A) and in the time bins representing distinct therapy phases as pseudotime (B); dashed vertical lines represent the transition timepoints between clinical phases in days relative to stem cell transplantation (A) and in pseudotime (B). Related to **Methods**.

**Figure S12. Different TaxUMAP parameter choices**. Related to **Methods**. In addition to sample-by-sample distances based upon ASV level differences (naïve UMAP), the indicated taxonomic level was included in the sample-by-sample distance estimation (top row). Multiple levels may be combined; in the main text, we combined Phylum and Family with equal weights (bottom row, left). Increased weights for Phylum (middle) or Family (right) accentuate corresponding compositional features of the measured microbial compositions in a sample.

**Table S1. The list of all ASV-3-*E. coli* and ASV-36-*Klebsiella* isolates used in this study and their NCBI accession numbers (“Accession Isolate” column)**. Related to **Figure 4** and **Figure S9**.

**Table S2. Total number of ASV-36 and ASV-3 colonies counted in each well of BIOLOG PM1 plate after 24 h incubation inside of anaerobic chamber**. Related to **Figure 5A**. Carbon sources are listed in the same order as in Figure 5A.

**Table S3. Total number of ASV-36 and ASV-3 colonies counted in each well of BIOLOG PM1 plate after 24 h incubation in aerobiosis**. Related to **Figure 5B**. Carbon sources are listed in the same order as in Figure 5B.

**Table S4. Antibiotic administration exposure in three clinical phases**. Related to **Methods**.

