## Supplementary figures and images for "The TaxUMAP atlas: efficient display of large clinical microbiome data reveals ecological competition involved in protection against bacteremia"

### Supplemental Figure 1

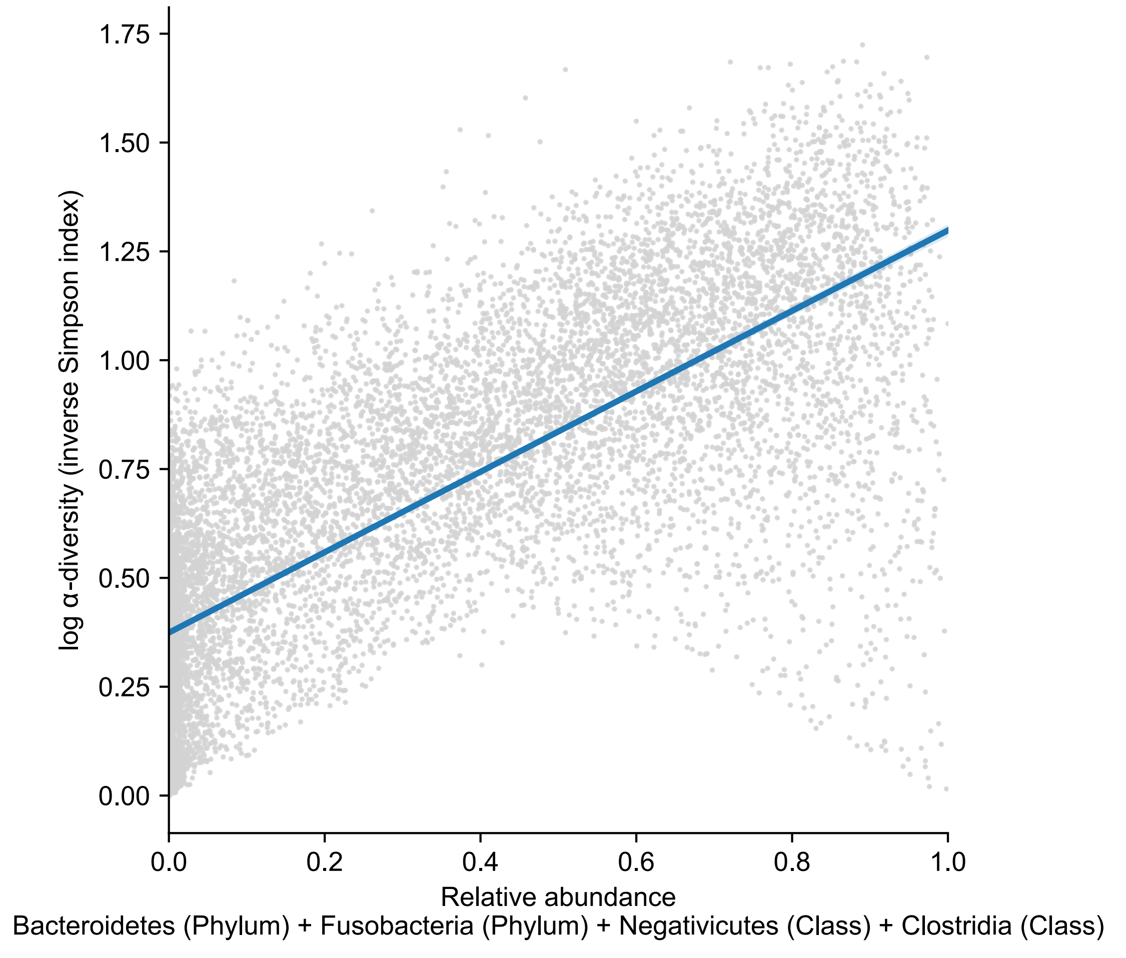

### Supplemental Figure 2

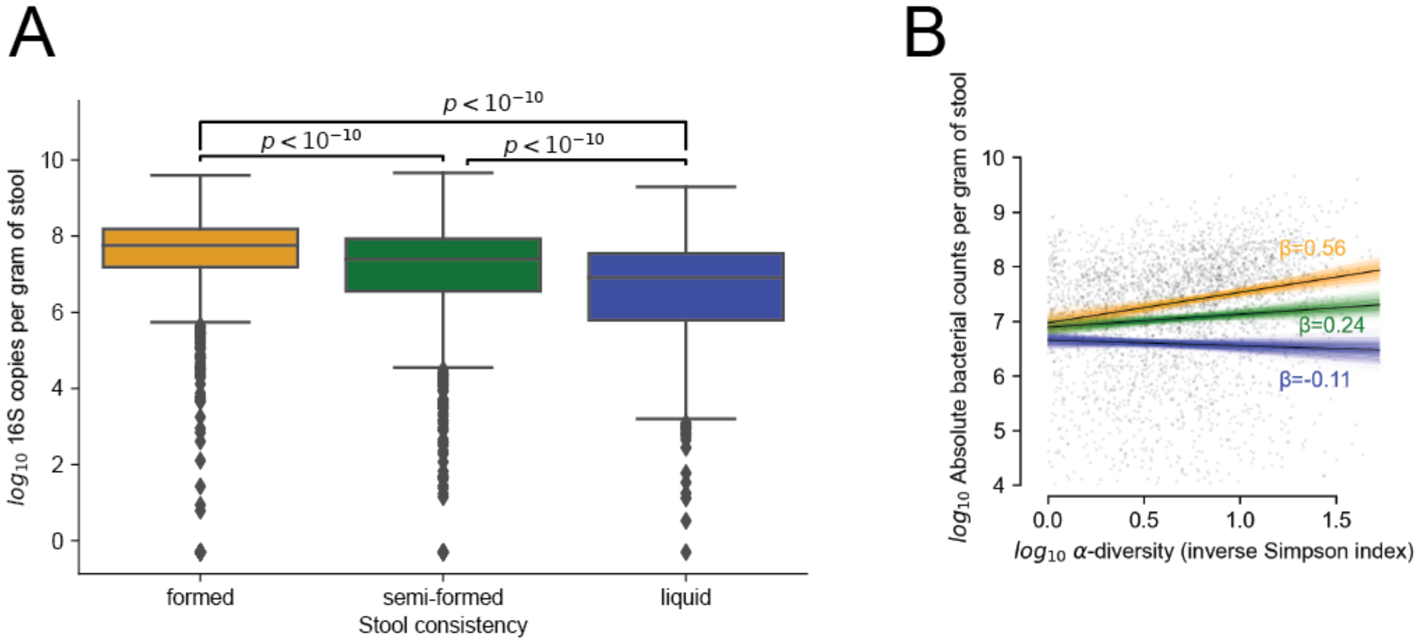

### Supplemental Figure 3

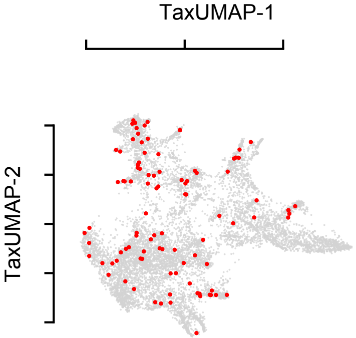

### Supplemental Figure 4

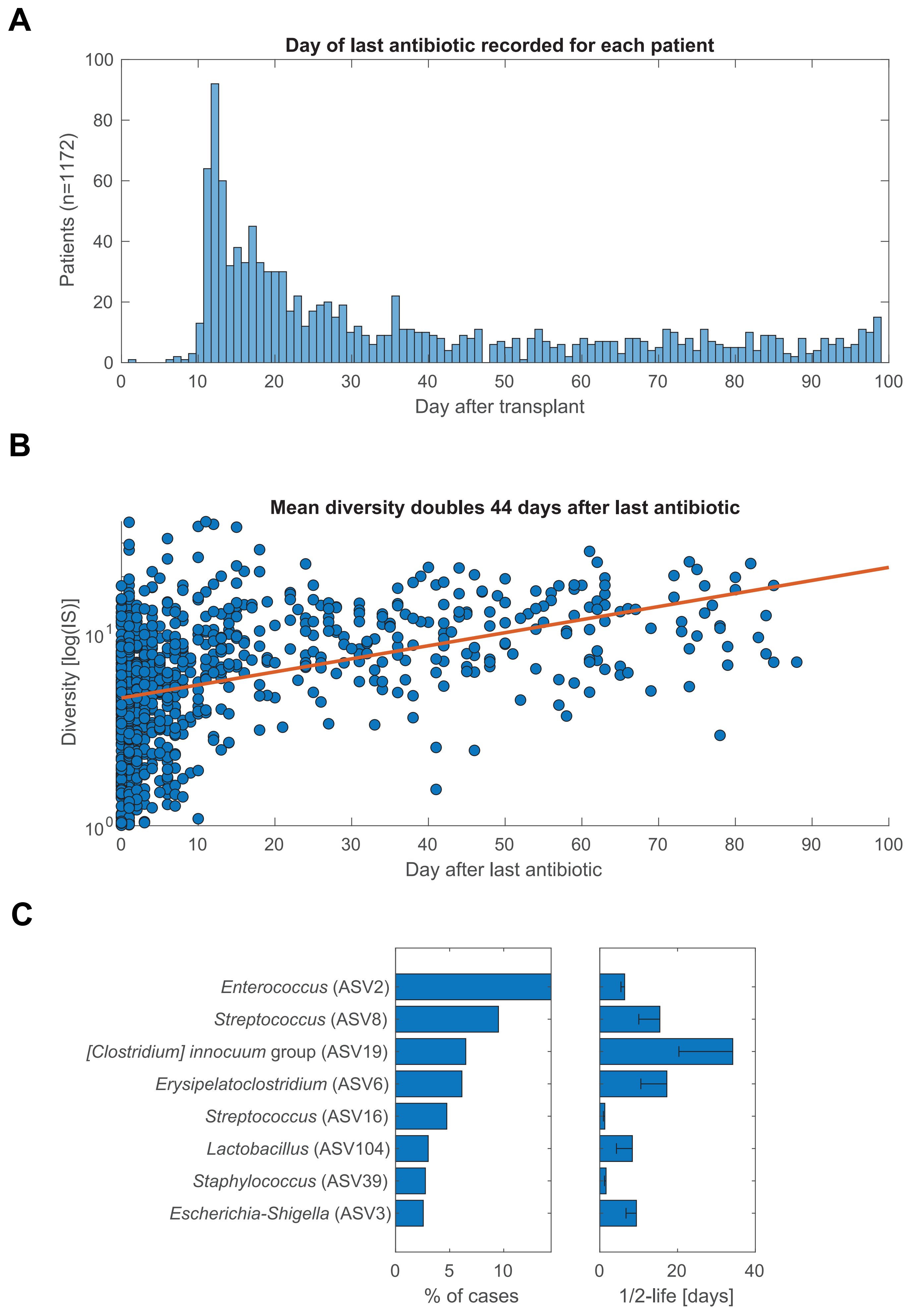

### Supplemental Figure 5

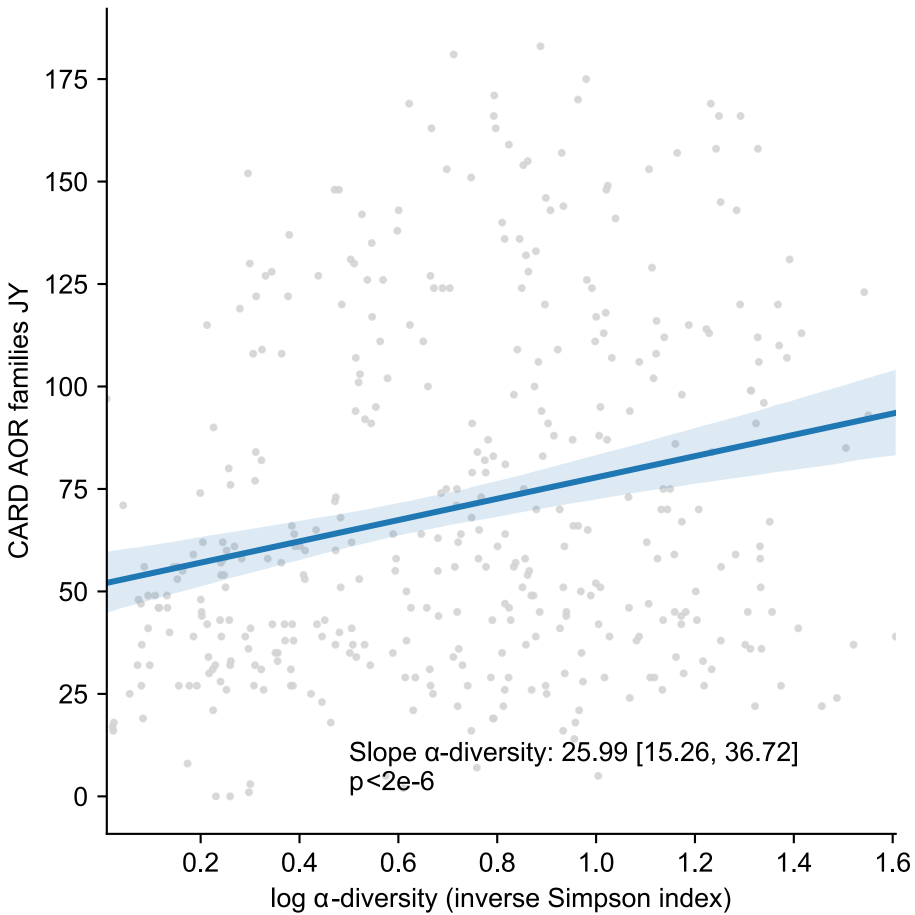

### Supplemental Figure 6

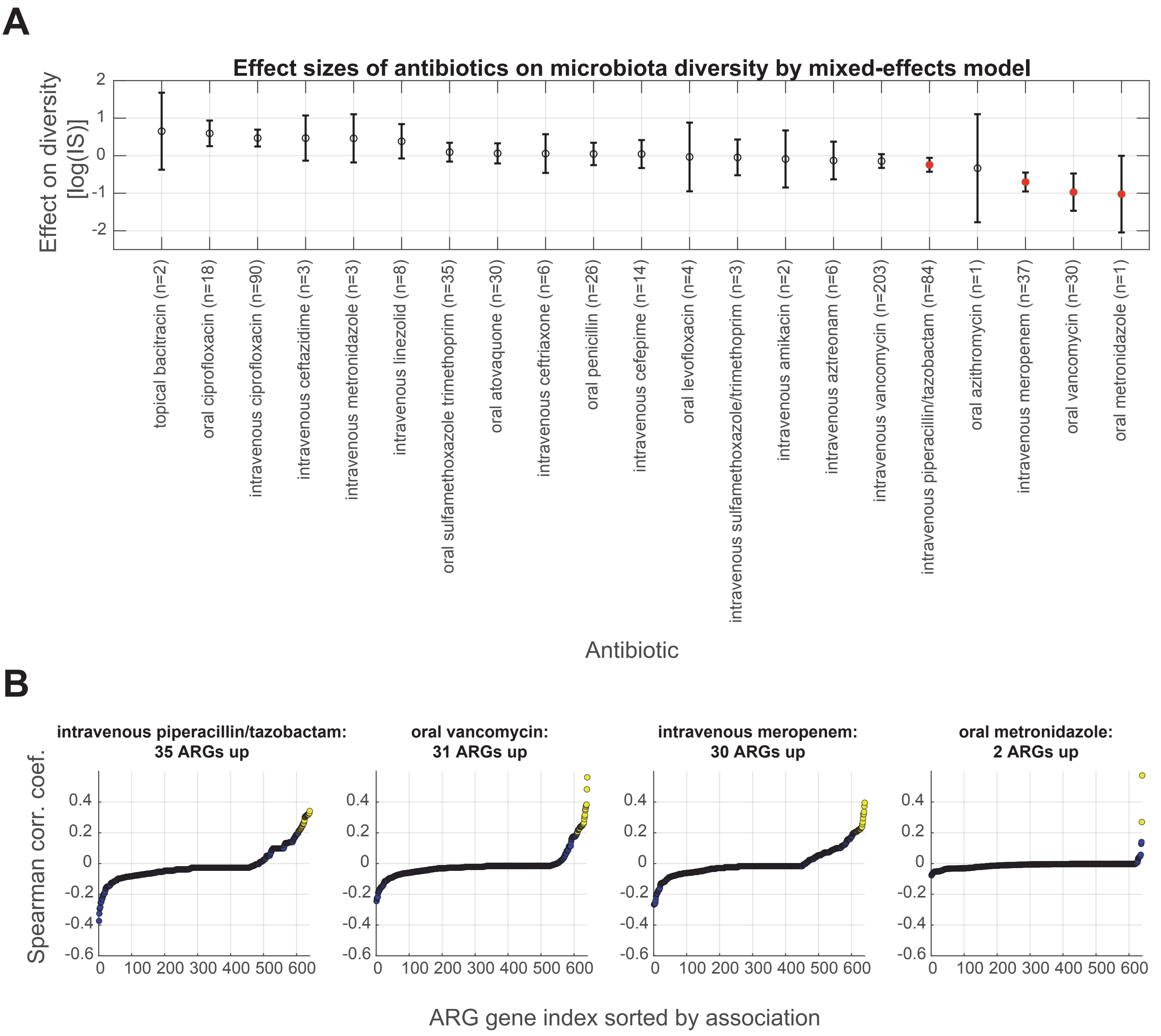

### Supplemental Figure 7

***vanA* gene detected in our patient cohort matches *vanA* gene from *E. faecium***

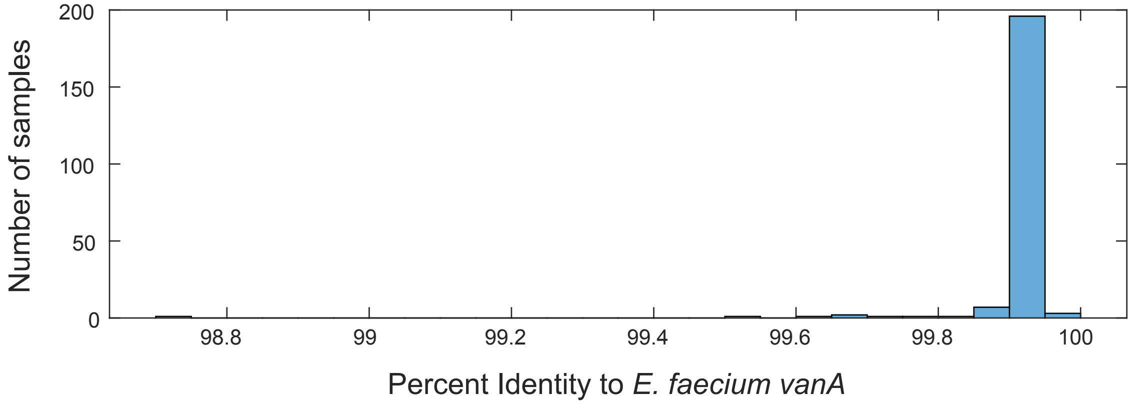

### Supplemental Figure 8

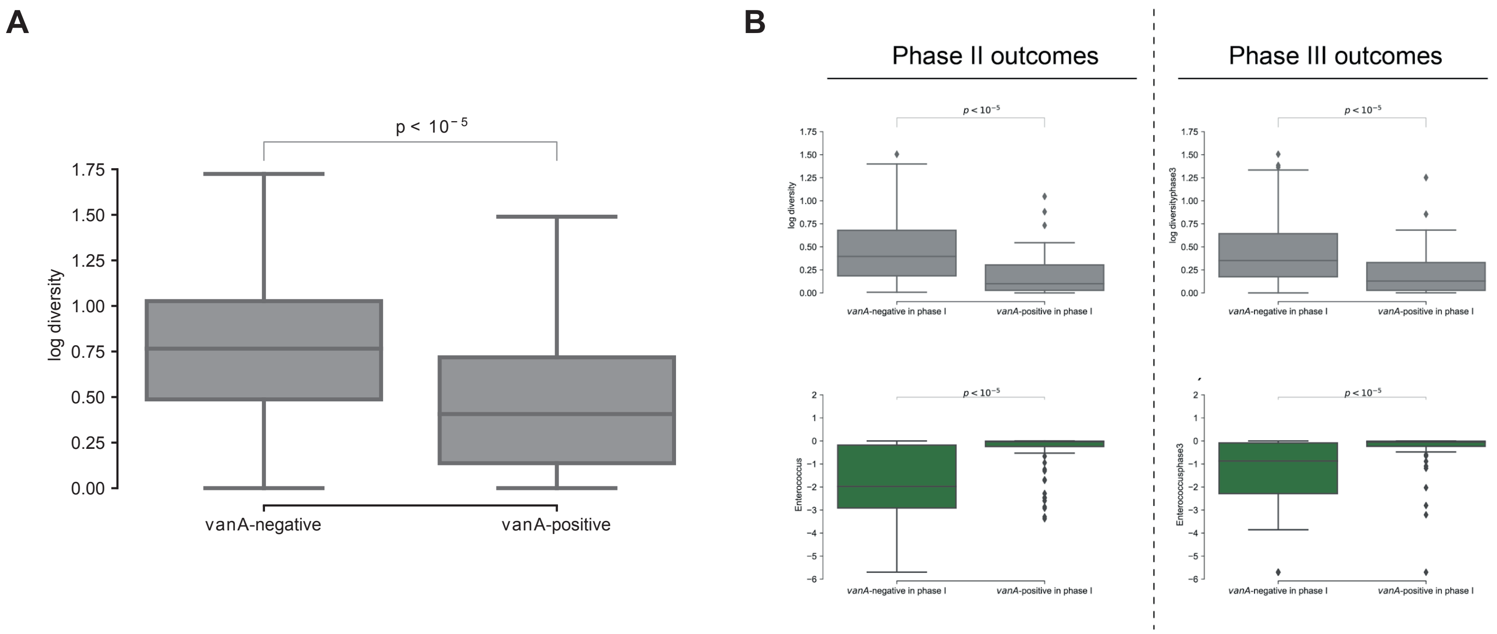

### Supplemental Figure 9

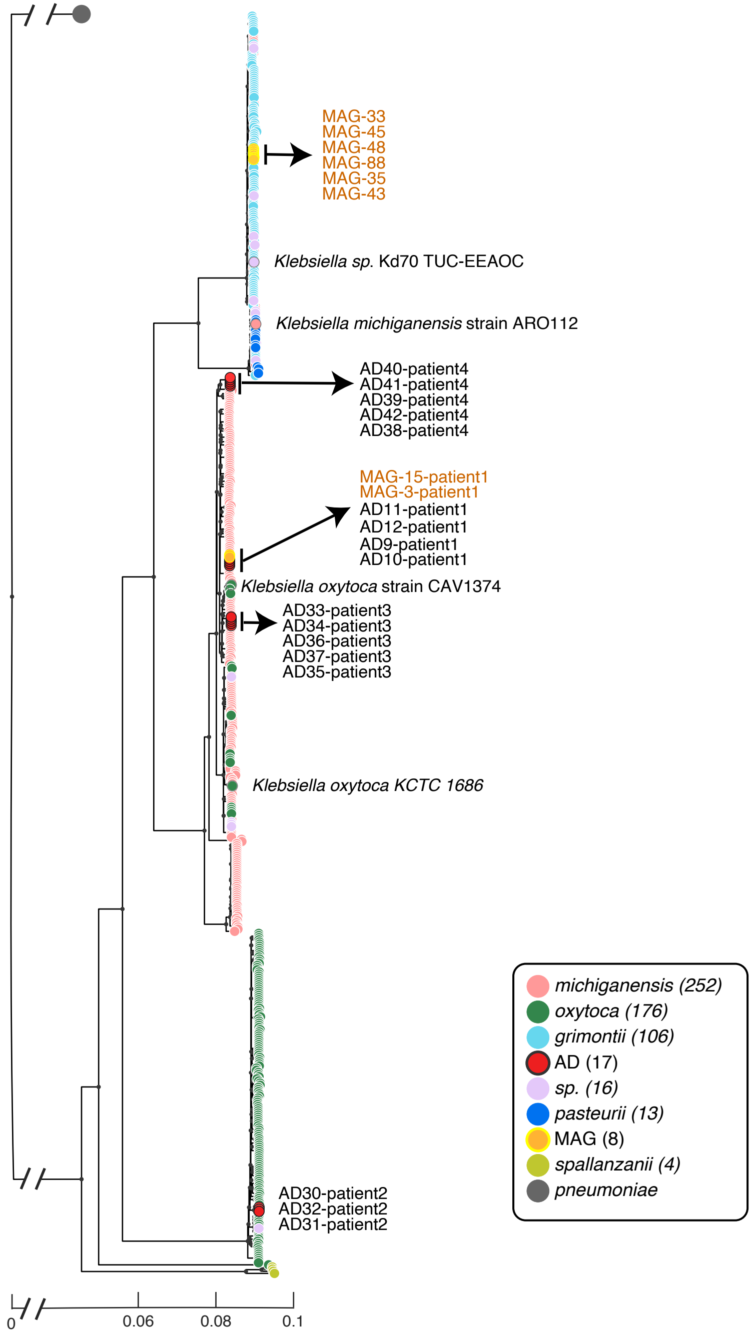

### Supplemental Figure 10

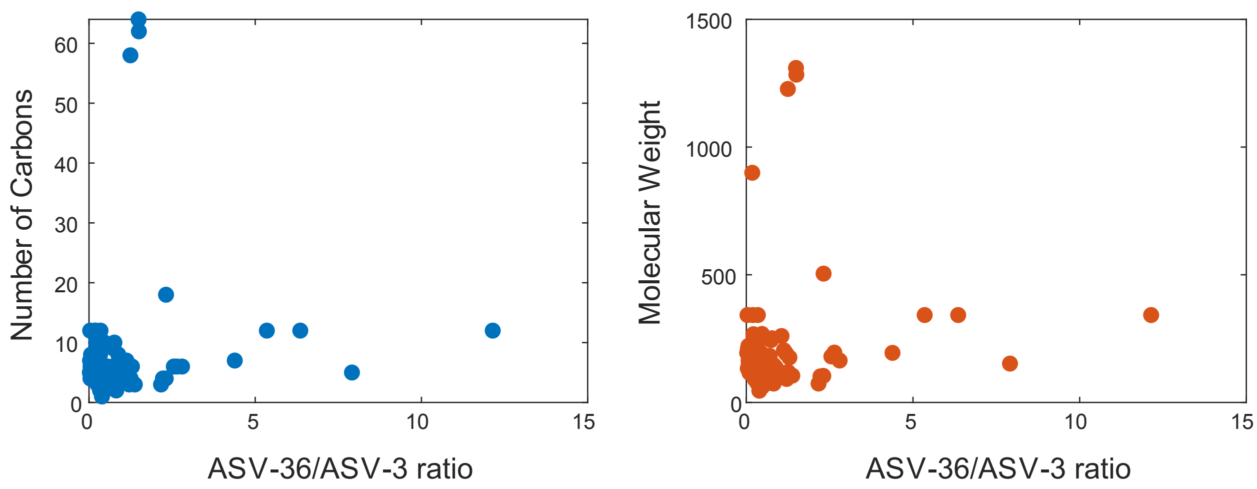

### Supplemental Figure 11

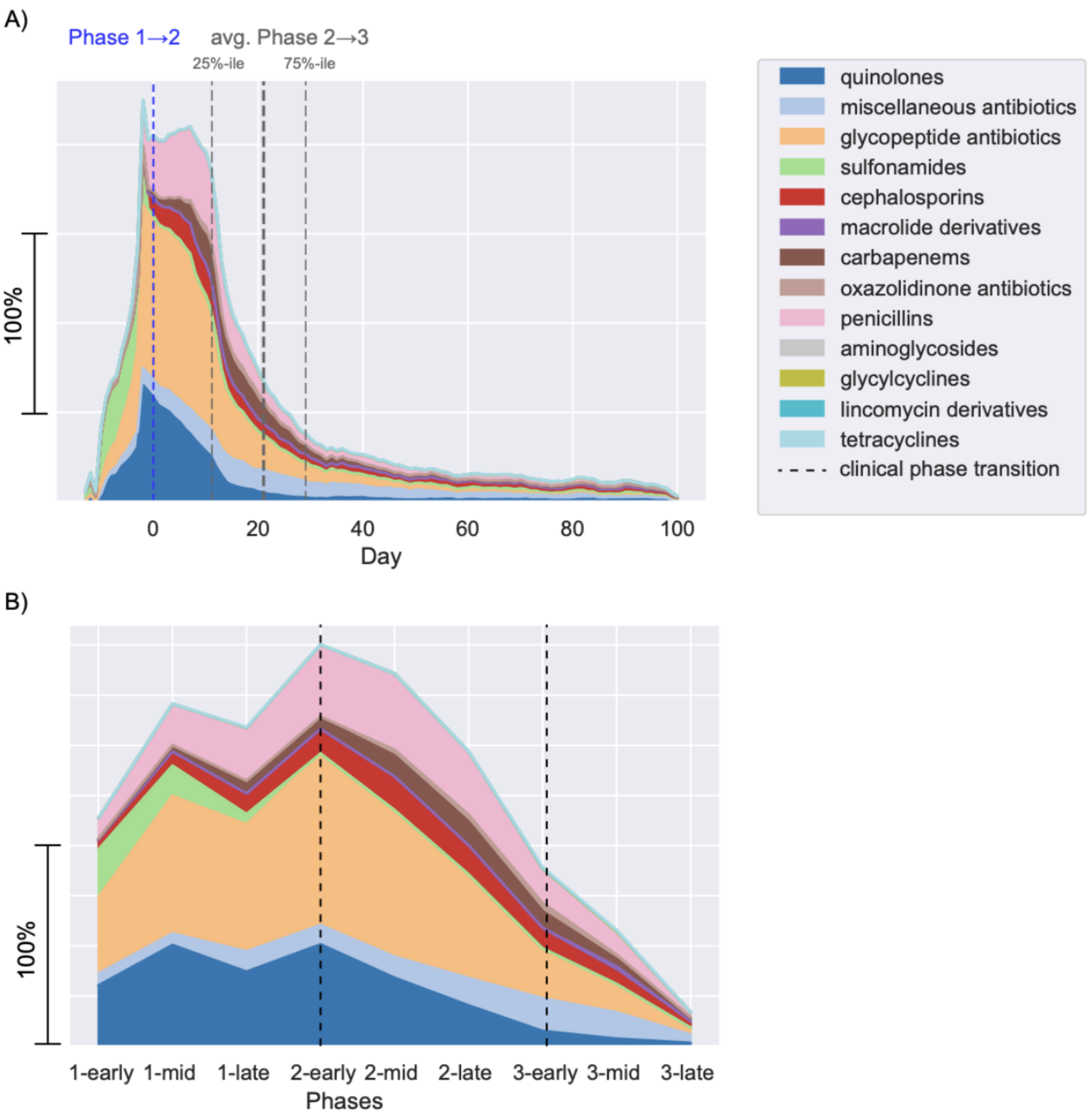

### Supplemental Figure 12

by ASV level differences and:

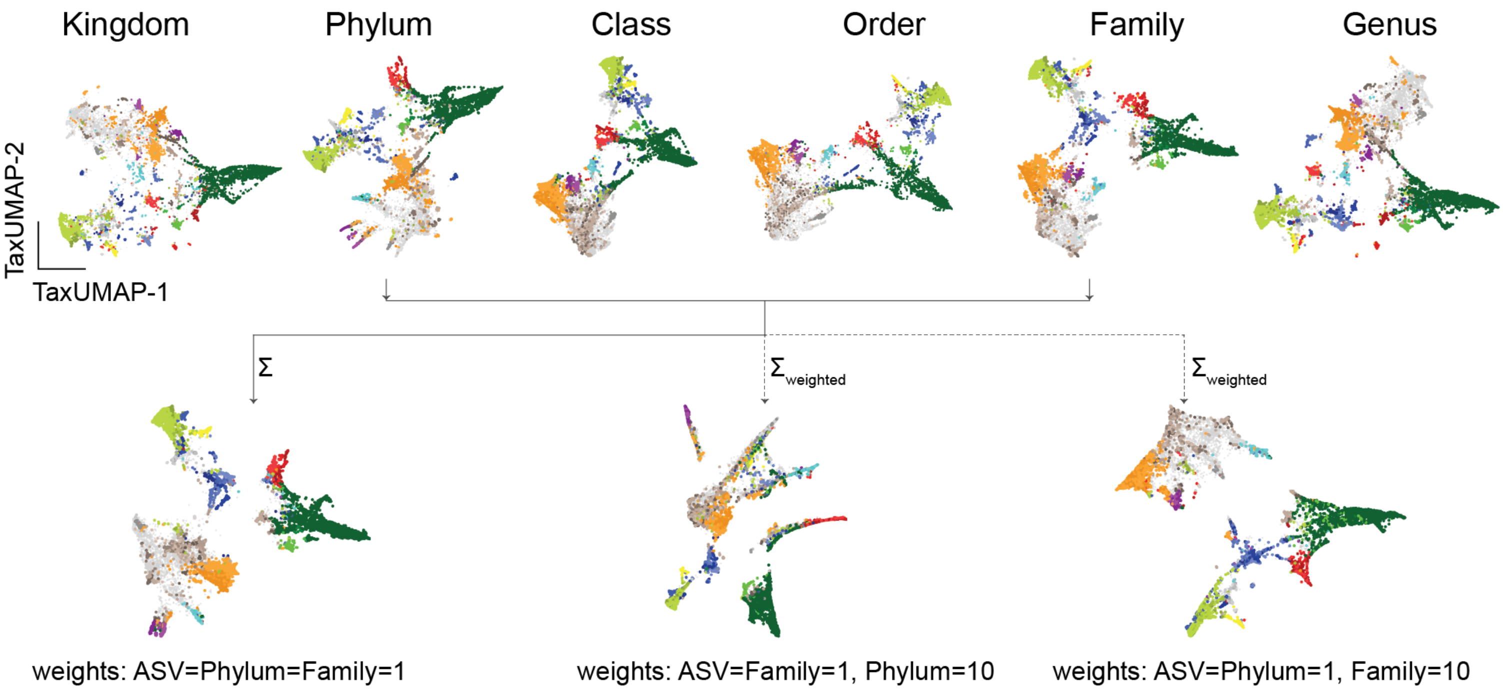
