## Supplemental Table 1 for "The TaxUMAP atlas: efficient display of large clinical microbiome data reveals ecological competition involved in protection against bacteremia"

| Isolate ID (as in Figure 4 and Figure S6) | ASV | Sample ID | Patient ID | BioProject | Accession Isolate |
|---|---|---|---|---|---|
| 9 | ASV-36-*Klebsiella* | FMT.0009U | FMT.0009 | PRJNA545312 | SRR14131380 |
| 10 | ASV-36-*Klebsiella* | FMT.0009U | FMT.0009 | PRJNA545312 | SRR14131379 |
| 11 | ASV-36-*Klebsiella* | FMT.0009U | FMT.0009 | PRJNA545312 | SRR14131378 |
| 12 | ASV-36-*Klebsiella* | FMT.0009U | FMT.0009 | PRJNA545312 | SRR14131376 |
| 24 | ASV-3-*E. coli* | 745A | 745 | PRJNA606262 | SRR14131383 |
| 25 | ASV-3-*E. coli* | 745A | 745 | PRJNA606262 | SRR14131382 |
| 26 | ASV-3-*E. coli* | 745A | 745 | PRJNA606262 | SRR14131381 |
| 30 | ASV-36-*Klebsiella* | 149D | 149 | PRJNA545312 | SRR14131374 |
| 31 | ASV-36-*Klebsiella* | 149D | 149 | PRJNA545312 | SRR14131373 |
| 32 | ASV-36-*Klebsiella* | 149D | 149 | PRJNA545312 | SRR14131372 |
| 33 | ASV-36-*Klebsiella* | 729E | 729 | PRJNA545312 | SRR14131389 |
| 34 | ASV-36-*Klebsiella* | 729E | 729 | PRJNA545312 | SRR14131387 |
| 35 | ASV-36-*Klebsiella* | 729E | 729 | PRJNA545312 | SRR14131386 |
| 36 | ASV-36-*Klebsiella* | 729E | 729 | PRJNA545312 | SRR14131385 |
| 37 | ASV-36-*Klebsiella* | 729E | 729 | PRJNA545312 | SRR14131384 |
| 38 | ASV-36-*Klebsiella* | 1392M | 1392 | PRJNA545312 | SRR14131400 |
| 39 | ASV-36-*Klebsiella* | 1392M | 1392 | PRJNA545312 | SRR14131399 |
| 40 | ASV-36-*Klebsiella* | 1392M | 1392 | PRJNA545312 | SRR14131388 |
| 41 | ASV-36-*Klebsiella* | 1392M | 1392 | PRJNA545312 | SRR14131377 |
| 42 | ASV-36-*Klebsiella* | 1392M | 1392 | PRJNA545312 | SRR14131375 |
| 48 | ASV-3-*E. coli* | 1814W | 1814 | PRJNA607574 | SRR16077561 |
| 49 | ASV-3-*E. coli* | 1814W | 1814 | PRJNA607574 | SRR16077560 |

**Table S1. The list of all ASV-3-*E. coli* and ASV-36-*Klebsiella* isolates used in this study and their NCBI accession numbers ("Accession Isolate" column). Related to Figure 4 and Figure S6.**
