## Supplemental Table 2 for "The TaxUMAP atlas: efficient display of large clinical microbiome data reveals ecological competition involved in protection against bacteremia"

| Metabolite | Average_CFUs |
|---|---|
| Inoculum | 1380000 |
| L_Lyxose | 11300000 |
| a_Methyl_D_Galactoside | 27866666.7 |
| a_D_Lactose | 39833333.3 |
| L_AsparticAcid | 5166666.67 |
| D_SaccharicAcid | 7733333.33 |
| L_Asparagine | 6900000 |
| L_MalicAcid | 11900000 |
| N_Acetyl_D_Glucosamine | 31566666.7 |
| L_Fucose | 32400000 |
| L_Arabinose | 23233333.3 |
| FumaricAcid | 21200000 |
| D_AsparticAcid | 7133333.33 |
| D_L_MalicAcid | 4833333.33 |
| MucicAcid | 19366666.7 |
| BromoSuccinicAcid | 7966666.67 |
| D_Sorbitol | 21200000 |
| D_Fructose | 18800000 |
| L_GlutamicAcid | 2300000 |
| M_TartaricAcid | 800000 |
| D_Mannitol | 33166666.7 |
| N_Acetyl_b_D_Mannosamine | 4000000 |
| D_Galactose | 11533333.3 |
| D_Melibiose | 24100000 |
| D_Trehalose | 23066666.7 |
| SuccinicAcid | 1933333.33 |
| Adenosine | 13133333.3 |
| Fructose_6_Phosphate | 16866666.7 |
| D_GlucuronicAcid | 12533333.3 |
| D_Ribose | 9233333.33 |
| Glycyl_L_Proline | 1633333.33 |
| MonoMethylSuccinate | 2500000 |
| P_HydroxyPhenylAceticAcid | 1300000 |
| Glucose_1_Phosphate | 26266666.7 |
| L_GalactonicAcid_g_Lactone | 11066666.7 |
| L_LacticAcid | 1600000 |
| PyruvicAcid | 10666666.7 |
| D_GluconicAcid | 10600000 |
| D_GalacturonicAcid | 14933333.3 |
| Glycyl_L_AsparticAcid | 5700000 |
| D_Xylose | 12933333.3 |

| | |
|---|---|
| L_Alanine | 3833333.33 |
| M_HydroxyPhenylAceticAcid | 1266666.67 |
| L_Proline | 1800000 |
| GlycolicAcid | 1800000 |
| Phenylethyl_amine | 366666.667 |
| Lactulose | 3433333.33 |
| a_D_Glucose | 15733333.3 |
| D_GalactonicAcid_g_Lactone | 9333333.33 |
| FormicAcid | 1966666.67 |
| Thymidine | 10066666.7 |
| AceticAcid | 1100000 |
| Glucuronamide | 5866666.67 |
| Inosine | 23266666.7 |
| Two_Aminoethanol | 1533333.33 |
| D_Mannose | 21833333.3 |
| Acetoaceticacid | 733333.333 |
| D_L_a_Glycerol_Phosphate | 4466666.67 |
| L_Alanyl_Glycine | 2000000 |
| D_Psicose | 2033333.33 |
| MethylPyruvate | 5600000 |
| NegativeControl | 1400000 |
| 1_2_Propanediol | 1800000 |
| a_Keto_GlutaricAcid | 2666666.67 |
| Dulcitol | 10266666.7 |
| Uridine | 17966666.7 |
| L_Serine | 7300000 |
| Two_DeoxyAdenosine | 3500000 |
| D_Alanine | 500000 |
| a_HydroxyGlutaricAcid_g_Lactone | 2200000 |
| D_MalicAcid | 2166666.67 |
| GlyoxylicAcid | 2366666.67 |
| L_Glutamine | 2100000 |
| Tyramine | 1533333.33 |
| D_Threonine | 400000 |
| Glucose_6_Phosphate | 16900000 |
| Glycyl_L_GlutamicAcid | 1500000 |
| CitricAcid | 3933333.33 |
| Glycerol | 5333333.33 |
| Tween20 | 2433333.33 |
| L_Threonine | 1500000 |
| TricarballylicAcid | 2666666.67 |
| D_Serine | 2266666.67 |

| | |
|---|---|
| Tween80 | 4200000 |
| Tween40 | 3100000 |
| Propionicacid | 700000 |
| a_Keto_ButyricAcid | 666666.667 |
| a_HydroxyButyricacid | 1333333.33 |
| Maltotriose | 4033333.33 |
| M_Inositol | 7300000 |
| D_GlucosaminicAcid | 3566666.67 |
| L_Rhamnose | 9766666.67 |
| b_Methyl_D_Glucoside | 6733333.33 |
| Sucrose | 9900000 |
| Maltose | 9033333.33 |
| Adonitol | 6666666.67 |
| D_Cellobiose | 12366666.7 |
