## Supplemental Table 3 for "The TaxUMAP atlas: efficient display of large clinical microbiome data reveals ecological competition involved in protection against bacteremia"

| Metabolite | Average_CFUs |
|---|---|
| Inoculum | 2716666.67 |
| Adenosine | 98666666.7 |
| Tyramine | 7333333.33 |
| a_Keto_ButyricAcid | 9333333.33 |
| N_Acetyl_b_D_Mannosamine | 21000000 |
| BromoSuccinicAcid | 81333333.3 |
| Glycyl_L_AsparticAcid | 78666666.7 |
| Propionicacid | 62000000 |
| Glucose_6_Phosphate | 104333333 |
| Glycyl_L_GlutamicAcid | 41000000 |
| SuccinicAcid | 77333333.3 |
| D_Psicose | 17000000 |
| D_L_MalicAcid | 152000000 |
| L_Lyxose | 95666666.7 |
| FumaricAcid | 106000000 |
| Acetoaceticacid | 7000000 |
| D_MalicAcid | 129666667 |
| Two_DeoxyAdenosine | 126333333 |
| M_TartaricAcid | 50666666.7 |
| Inosine | 158333333 |
| a_D_Lactose | 244000000 |
| Phenylethyl_amine | 6666666.67 |
| a_Methyl_D_Galactoside | 209000000 |
| Glucose_1_Phosphate | 165333333 |
| L_MalicAcid | 110666667 |
| D_Galactose | 110333333 |
| Uridine | 213666667 |
| D_SaccharicAcid | 94333333.3 |
| L_LacticAcid | 114666667 |
| D_GalacturonicAcid | 123666667 |
| D_GlucuronicAcid | 137666667 |
| D_Threonine | 0 |
| Fructose_6_Phosphate | 111666667 |
| L_GalactonicAcid_g_Lactone | 59666666.7 |
| AceticAcid | 53666666.7 |
| PyruvicAcid | 82333333.3 |
| D_GluconicAcid | 131666667 |
| D_Trehalose | 216666667 |
| D_Sorbitol | 257666667 |
| a_HydroxyButyricacid | 2000000 |
| a_D_Glucose | 248000000 |

| | |
|---|---|
| D_Alanine | 218000000 |
| D_Mannitol | 274666667 |
| N_Acetyl_D_Glucosamine | 259666667 |
| L_AsparticAcid | 125333333 |
| D_Melibiose | 202333333 |
| L_Arabinose | 191666667 |
| D_Xylose | 247333333 |
| L_Fucose | 183000000 |
| L_Serine | 87666666.7 |
| D_Mannose | 246666667 |
| a_Keto_GlutaricAcid | 84666666.7 |
| D_Fructose | 247000000 |
| D_GalactonicAcid_g_Lactone | 171333333 |
| MonoMethylSuccinate | 14333333.3 |
| D_Ribose | 210666667 |
| Glycerol | 212333333 |
| D_AsparticAcid | 20000000 |
| Glycyl_L_Proline | 127666667 |
| MethylPyruvate | 47000000 |
| Tween80 | 16333333.3 |
| L_Alanyl_Glycine | 201666667 |
| Glucuronamide | 26000000 |
| L_Alanine | 188000000 |
| Two_Aminoethanol | 7333333.33 |
| NegativeControl | 22000000 |
| Thymidine | 142666667 |
| MucicAcid | 188666667 |
| Tween40 | 27666666.7 |
| GlyoxylicAcid | 178000000 |
| L_Rhamnose | 199666667 |
| 1_2_Propanediol | 14666666.7 |
| Tween20 | 27666666.7 |
| a_HydroxyGlutaricAcid_g_Lactone | 7000000 |
| b_Methyl_D_Glucoside | 272333333 |
| L_Asparagine | 59000000 |
| L_GlutamicAcid | 122000000 |
| L_Glutamine | 122333333 |
| D_L_a_Glycerol_Phosphate | 52666666.7 |
| M_Inositol | 162666667 |
| GlycolicAcid | 221666667 |
| Sucrose | 300333333 |
| D_Serine | 143333333 |

| | |
|---|---|
| D_Cellobiose | 240000000 |
| Maltotriose | 359000000 |
| Maltose | 249666667 |
| CitricAcid | 82666666.7 |
| P_HydroxyPhenylAceticAcid | 133333333 |
| D_GlucosaminicAcid | 146000000 |
| L_Threonine | 77000000 |
| M_HydroxyPhenylAceticAcid | 138000000 |
| L_Proline | 286000000 |
| FormicAcid | 47000000 |
| TricarballylicAcid | 126000000 |
| Dulcitol | 343333333 |
| Adonitol | 530000000 |
| Lactulose | 469000000 |
