## Supplemental Table 4 for "The TaxUMAP atlas: efficient display of large clinical microbiome data reveals ecological competition involved in protection against bacteremia"

| | | Phase | | |
|---|---|---|---|---|
| antibiotic category | days administered per patient | 1 | 2 | 3 |
| aminoglycosides | mean | 0.050524 | 0.108803 | 0.028634 |
| | std | 0.597643 | 0.803477 | 0.246822 |
| carbapenems | mean | 0.876072 | 2.115727 | 1.572687 |
| | std | 3.312291 | 4.971947 | 4.643664 |
| cephalosporins | mean | 1.177312 | 2.143422 | 1.829295 |
| | std | 3.236395 | 4.415567 | 4.722806 |
| glycopeptide antibiotics | mean | 7.711153 | 10.69535 | 4.044053 |
| | std | 6.305629 | 7.476178 | 6.725049 |
| glycylcyclines | mean | 0.026692 | 0.038576 | 0.024229 |
| | std | 0.501194 | 0.522272 | 0.524449 |
| lincomycin derivatives | mean | 0.055291 | 0.016815 | 0.085903 |
| | std | 1.3123 | 0.260833 | 1.393519 |
| macrolide derivatives | mean | 0.171592 | 0.271019 | 1.068282 |
| | std | 1.319667 | 2.046427 | 4.616705 |
| miscellaneous antibiotics | mean | 1.734986 | 2.6182 | 4.394273 |
| | std | 4.631901 | 5.221493 | 8.721392 |
| oxazolidinone antibiotics | mean | 0.222116 | 0.723046 | 0.632159 |
| | std | 1.385326 | 2.749699 | 2.432131 |
| penicillins | mean | 3.062917 | 5.443126 | 3.114537 |
| | std | 4.952816 | 6.662997 | 6.33397 |
| quinolones | mean | 4.125834 | 4.757666 | 1.39207 |
| | std | 4.220699 | 6.005907 | 3.378123 |
| sulfonamides | mean | 1.35939 | 0.09001 | 0.375551 |
| | std | 2.539292 | 1.028702 | 2.250796 |
| tetracyclines | mean | 0.019066 | 0.007913 | 0.035242 |
| | std | 0.407022 | 0.160249 | 0.410545 |
